# A phased, diploid assembly of the Cascade hop *(Humulus lupulus)* genome reveals patterns of selection and haplotype variation

**DOI:** 10.1101/786145

**Authors:** Lillian K. Padgitt-Cobb, Sarah B. Kingan, Jackson Wells, Justin Elser, Brent Kronmiller, Daniel Moore, Gregory Concepcion, Paul Peluso, David Rank, Pankaj Jaiswal, John Henning, David A. Hendrix

## Abstract

Hop *(Humulus lupulus* L. var Lupulus) is a diploid, dioecious plant with a history of cultivation spanning more than one thousand years. Hop cones are valued for their use in brewing, and around the world, hop has been used in traditional medicine to treat a variety of ailments. Efforts to determine how biochemical pathways responsible for desirable traits are regulated have been challenged by the large, repetitive, and heterozygous genome of hop. We present the first report of a haplotype-phased assembly of a large plant genome. Our assembly and annotation of the Cascade cultivar genome is the most extensive to date. PacBio long-read sequences from hop were assembled with FALCON and phased with FALCON-Unzip. Using the diploid assembly to assess haplotype variation, we discovered genes under positive selection enriched for stress-response, growth, and flowering functions. Comparative analysis of haplotypes provides insight into large-scale structural variation and the selective pressures that have driven hop evolution. Previous studies estimated repeat content at around 60%. With improved resolution of long terminal retrotransposons (LTRs) due to long-read sequencing, we found that hop is nearly 78% repetitive. Our quantification of repeat content provides context for the size of the hop genome, and supports the hypothesis of whole genome duplication (WGD), rather than expansion due to LTRs. With our more complete assembly, we have identified a homolog of cannabidiolic acid synthase (CBDAS) that is expressed in multiple tissues. The approaches we developed to analyze a phased, diploid assembly serve to deepen our understanding of the genomic landscape of hop and may have broader applicability to the study of other large, complex genomes.

## Background

Hop *(Humulus lupulus* L. var Lupulus) is a diploid (2n=18+XX/XY) [1], dioecious plant with a history of cultivation spanning over one thousand years [2–4]. Large size (2.7Gb), abundant repeat content, and heterozygosity have challenged draft assemblies of the hop genome [5, 6]. Hop cones, the flowers from the female plant, are valued for their use in brewing, and hop has been used worldwide in traditional medicine to treat a variety of ailments, including anxiety, insomnia, and pain. Modern research has revealed numerous compounds of medicinal interest in hop with activity against metabolic syndrome [7–9], and with anti-cancer [2, 10, 11], anti-microbial [2, 8, 12], and phytoestrogenic [13–16] properties. Cascade is a backcross hybrid of English and Russian varieties of hop [17] and is the most widely used hop cultivar in craft brewing [18].

Hop cones are rich in secondary metabolites, including bitter acids, terpenes, and polyphenols [8, 19, 20]. Hop was originally added to beer for its preservative activity and flavor, due to the bitter acids found in hop cones, including alpha and beta acids. Alpha acids, including humulone, cohumulone, and adhumulone, possess antimicrobial activity and contribute to beer foam stability [2, 21, 22]. Isomerization of alpha acids to iso-alpha-acids at high pH and temperature results in a characteristic bitter flavor [2]. The most abundant terpenes are monoterpene beta-myrcene, as well as sesquiterpenes alpha-humulene and beta-caryophyllene [23]. Hop also contains a variety of flavonoids [24], which are known for their health benefit [25]. The most abundant prenylflavonoid in hop is xanthohumol (XN), which features broad-spectrum anti-cancer activity. Hop is also known for containing 8-prenylnaringenin (8-PN), which has strong estrogenic activity and potential for pharmacological applications [13]. Through hop breeding, the development of new cultivars that are optimized to yield desirable pharmacologically-relevant metabolites would benefit from a more complete assembly, and the mapping of regulatory regions that direct genes involved in the biosynthesis of these compounds.

Powdery mildew reduces the quality and quantity of yield while increasing the cost of production [26]. Although Cascade has historically possessed disease resistance to fungal pathogens powdery mildew *(Podosphaero macularis)* [18] and downy mildew *(Pseudoperonospora humuli)* [27], there are still risks of damage by these fungal pathogens. A recent outbreak of powdery mildew affecting Cascade in the U.S. Pacific Northwest (PNW) was the result of a Cascade-adapted isolate of *P. Macularis* [18]. Very little management was required to grow Cascade until powdery mildew was introduced in the PNW in the mid-1990s [28], and the rise of PM infection corresponds to an increase in Cascade acreage [18]. Hop breeding with resistant cultivars [26, 27, 29] can result in new resistance, but given the inevitability of pathogen adaptation, an understanding of the genomic features underlying resistance and susceptibility is necessary to breed for hop that can withstand pathogen adaptations. Past efforts in the identification of alleles, genes, and regulatory genomic regions have been hindered by an incomplete draft assembly [26]. Insufficient genome coverage was cited as the reason for difficulty detecting an R-gene quantitative trait locus (QTL) for powdery mildew in hop [30]. Further work to identify QTLs associated with disease susceptibility and resistance will benefit from the updated, more-complete assembly, and will advance strategies for breeding disease-resistant hop.

Recent analyses of high-quality assemblies of cultivated plant species have revealed that genes and regulatory elements associated with desired traits are often located near long terminal retrotransposons (LTRs) [31–35]. LTRs are the most abundant type of transposable element (TE) in plant genomes and are largely responsible for the expansion of genome size, as they transpose by duplicating via a copy-and-paste mechanism [36]. Proliferation of TEs can occur as a response to biotic and abiotic stresses [37], and TE abundance in large plant genomes can be more than 80% [38]. TE proliferation is associated with a variety of changes to the genome, including gene regulation, duplication, rearrangement, and interruption of gene function [39]. In plant genomes, an average of 65% of genes are duplicated [40]. Most duplicated genes in plants are derived from whole genome duplication (WGD) events, transposon movement, and unequal crossing-over [40]. Duplicate genes provide a mechanism for plants to adapt to biotic and abiotic stresses, and are associated with speciation and diversity among plant species (40].

Most assemblies are collapsed, haploid representations of the genome, consisting of merged consensus sequences. Collapsed assemblies lose haplotype-specific information, confounding the identification of SNPs and structural variants. Regions of heterozygosity can also cause contig breakage, introducing more complexity into the assembly, and are particularly detrimental for assembling a large and highly heterozygous genome such as hop. Previous attempts at assembling the hop genome with short-reads have been fragmented and incomplete, containing collapsed regions that thwart the investigation of repeats, genes embedded within repeats, gene copy number, and heterozygosity (5, 6]. Previously, *H. lupulus* was estimated at ∼60% repetitive (42]. However, incompletely assembled and collapsed LTR regions obscured the true extent of repeat content in the genome. With long-read sequencing, the challenge of assembly fragmentation can be overcome [41]. PacBio single molecule, real-time sequencing (SMRT) produces reads spanning an average of 10-16 kb (43], up to nearly 100 kb (44]. In addition, algorithmic advances in long-read assembly have given rise to phased assemblies (45], in which both haplotypes are assembled independently. Haplotype-phased assemblies allow for investigation of genome structure beyond what is possible for a collapsed haploid assembly. Long-read sequencing enables a high-quality, more-complete assembly capable of resolving long terminal retrotransposons (LTRs), regions of high heterozygosity, and gene copy number variation.

## Results

The large size and complexity of the hop genome, a member of the Cannabaceae family (Figure 1a), has challenged previous assembly efforts. The goal of a phased diploid assembly is to create a set of primary contigs and associate contigs corresponding to the alternate haplotype. The haplotype-phased assembly of Cascade began with PacBio SMRT sequencing of DNA extracted from leaf tissue (Figure 1b). Assembly was performed with FALCON and phasing was performed with FALCON-Unzip (46], which separated the assembly into primary contigs and shorter associate contigs called “haplotigs.” The draft primary assembly was 4.24 Gb, containing 11,705 contigs, and the draft associate assembly was 1.35 Gb, containing 38,060 haplotigs. Low heterozygosity in the primary assembly resulted in collapsed primary contigs, where not enough variation was present to perform phasing. Medium heterozygosity, defined as less than 4% divergence between haplotypes, was optimal for phasing. In regions of high heterozygosity, primary contigs too diverged to be recognized as haplotypes were assembled as independent primary contigs. We further evaluated the primary assembly to identify duplicated primary contigs and reassign the shorter primary contig as a type of associate contig called a “homologous primary contig” (HPC) (Figure 1e).

**Figure 1.**
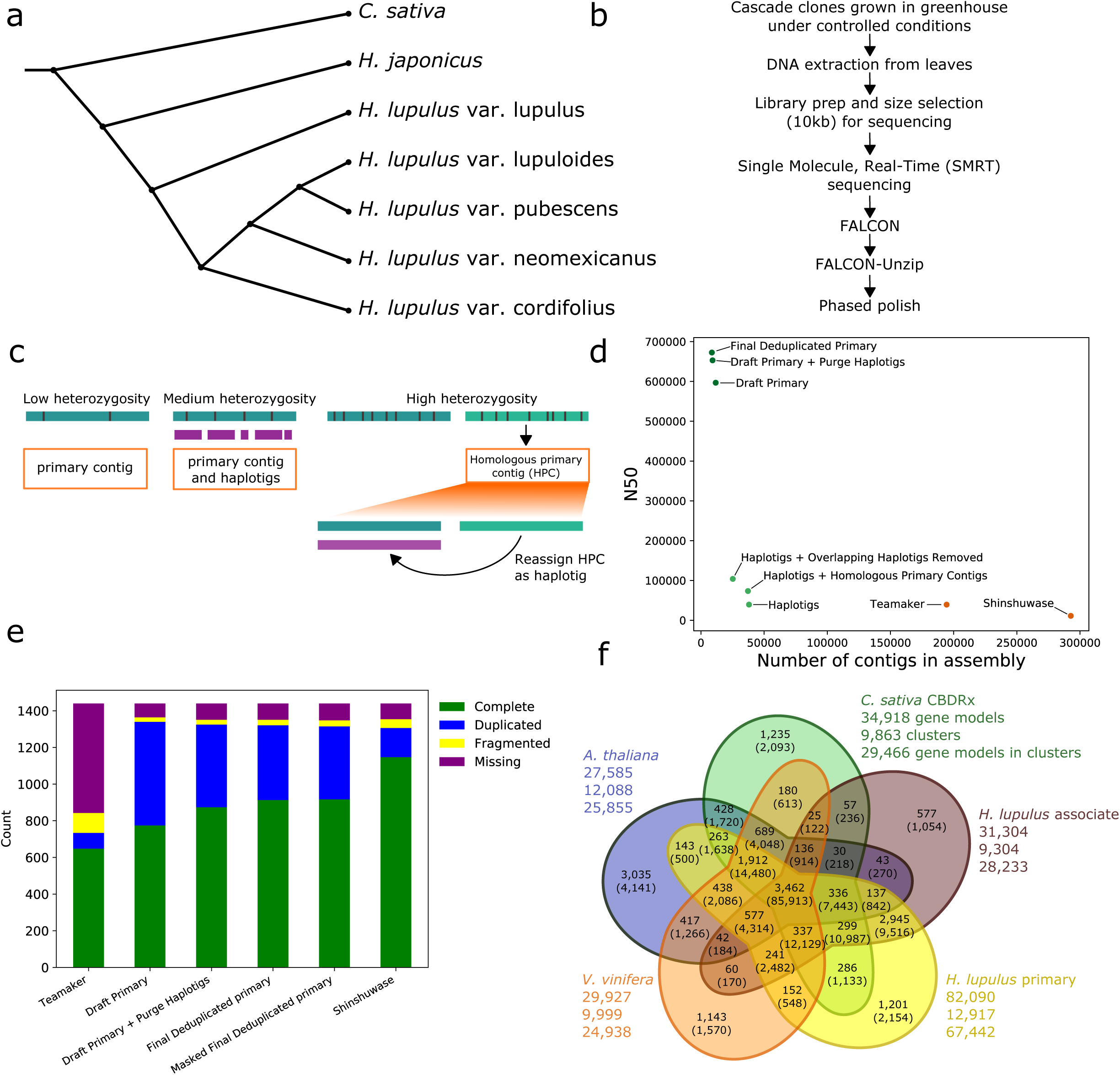

Multiple approaches were enlisted to perform deduplication. Although 2,491 of the 11,705 primary contigs were identified as HPCs by purge_haplotigs, the 3.82Gb primary assembly was still far larger than the estimated genome size (2.7Gb), and 451 duplicated BUSCO genes remained (874 single-copy; 451 duplicated). We developed a pairwise, sequence alignment-based approach to further identify and reassign HPCs. We identified an additional 568 HPCs, reducing the primary assembly to 3.71Gb (8,861 contigs) without excessive loss of complete, single-copy BUSCO genes (913 single-copy; 408 duplicated). The size of the haplotig assembly increased to 1.78 Gb (37,223 contigs). The N50 of the draft primary assembly was 596.74kb, and the N50 of the primary assembly following deduplication with purge_haplotigs was 652.97kb. Ultimately, the N50 of the final, deduplicated primary assembly was 672.6kb. The N50 of the draft haplotig assembly was 39.43kb, while the N50 of the haplotig assembly following reassignment of HPCs was 73.5kb (Figure 1d, Table 1). Many haplotigs and HPCs identified by purge_haplotigs overlapped in position on their corresponding primary contig. The shorter of the overlapping haplotigs and HPCs were removed from the haplotig assembly for downstream analyses. Following overlap removal, the N50 of the haplotig assembly was increased to 103.97kb. Haplotigs comprise 65.88% of the associate assembly and cover 26.5% of the primary assembly, while HPCs comprise 34.12% of the associate assembly and cover 13.73% of the primary assembly (Table 2).

**Table 1:** Assembly statistics

| Metric | Draft primary | purge_haplotigs | Deduplicated primary | Draft Associate | Associate (no overlap filtering) | Associate (overlaps filtered) | Shinshuwase | Teamaker |
| --- | --- | --- | --- | --- | --- | --- | --- | --- |
| Assembly length | 4.24Gb | 3.82Gb | 3.71Gb | 1.35Gb | 1.78Gb | 1.49Gb | 1.81Gb | 2.77 |
| Total number of contigs | 11705 | 9214 | 8661 | 38060 | 37223 | 25239 | 292698 | 194438 |
| N50 | 596.74kb | 652.97kb | 672.60kb | 39.43kb | 73.50kb | 103.97kb | 11.13kb | 39.33kb |
| Largest contig | 8.25Mb | 8.25Mb | 8.25Mb | 1.77Mb | 1.77Mb | 1.77Mb | 129.33kb | 367.72kb |

**Table 2.** Associate assembly percent coverage

| Percent assembly coverage | Deduplicated primary | Associate |
| --- | --- | --- |
| Haplotig (overlap unfiltered) | 33.7 | 70.3 |
| HPC (overlap unfiltered) | 14.24 | 29.7 |
| Haplotig (overlap filtered) | 26.5 | 65.88 |
| HPC (overlap filtered) | 13.73 | 34.12 |

The final deduplicated assembly was selected based on reduction of assembly size and best BUSCO results (Figure 1e, Supplementary Table 1). For BUSCO assessment, the goal was to maximize the number of single-copy complete genes, while minimizing the number of duplicated and missing BUSCO genes. The final deduplicated assembly recovered 1,321 out of 1,440 BUSCO genes (91.7%). Out of 1,321 complete BUSCO genes, 913 are single-copy (63.4%) and 408 are duplicated (28.3%). Although the Shinshuwase assembly contains more single-complete BUSCO genes (1,147) and fewer duplicated BUSCO genes (159), the assembly is highly fragmented, given the large number of scaffolds comprising the assembly (292,698), low N50 (11.13kb) (Figure 1d, Table 1), and short scaffold length (Supplementary Figure 1c). The relatively low repeat content of the Shinshuwase assembly suggests that scaffolds were broken in repeats and that repeats are collapsed in the assembled contigs. The fragmented composition of the short-read assemblies, as demonstrated by low N50 and large number of short scaffolds suggests they are composed of gene islands and do not comprehensively capture intergenic and transposon regions, as well as gene duplication events.

We assessed the association of protein-coding genes in the primary and haplotig assemblies with protein-coding genes from 115 species representing a range of clades from the tree of life (Figure 1f, Supplementary Table 2) clustered in 117,088 gene families. A comparison of 21,014 gene family clusters containing at least one of the genes from *A. thaliana, V. vinifera,* C. *sativa,* as well as the primary and haplotig assemblies, revealed that ∼go% of haplotig genes and 82% of genes from primary assembly have gene family assignments, as compared to 93% of *A. thaliana* and ∼84% of genes from C. *sativa* and *V. vinifera.* Between the five genome assemblies, there were 3,462 common gene families. We also found that 1,054 and 2,154 genes were unique to the haplotig and primary assemblies present in 577 and 1,201 gene families, respectively (Supplementary Table 3). At least half of these gene families contain more than one hop gene family member, while the rest are shared with genes from the remaining species.

We identified 1,679 non-redundant Gypsy, Copia, and unknown-type retrotransposons with a *de novo* approach, using LTR_FINDER [47], LTRharvest [48], and LTR_retriever [49]. We then combined the set of *de novo* LTR sequences with a repeat database from MIPS PlantDB [SO] and annotated the assembly with RepeatMasker [51]. We next assessed the repeat content of the deduplicated genome (Figure 2). We found that LTRs comprised 96.4% of total repeat content by length, which is over twice as much as *Arabidopsis,* and comparable to maize (Figure 2a). When examined as a proportion of the total genome length, the deduplicated assembly is nearly 78% repetitive (Figure 2b, Table 3). Gypsy-type elements are the longest type of LTR in hop, with an average length of 4,000 base pairs, and are also the most abundant (Figure 2c). Given the high repeat content of the hop genome, we investigated whether repeat content was responsible for the greater heterozygosity in primary contigs associated with HPCs compared to haplotigs. We found that both types of associate contig contain a similar percentage of repeat content (Figure 2d) and concluded that repeat content was not likely the source of divergence.

**Figure 2.**
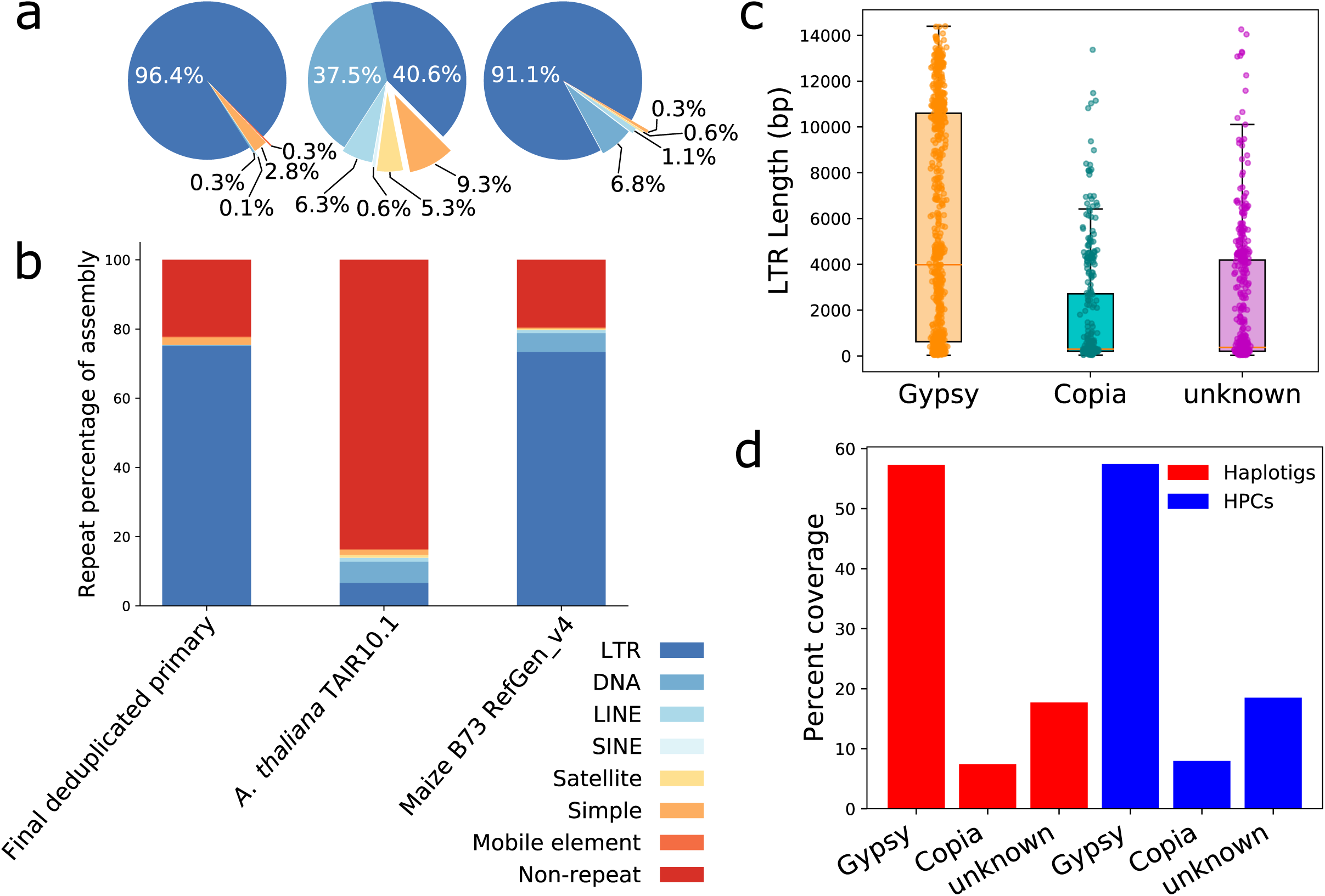

**Table 3:** Percent of assembly covered by repeats

| Category | Deduplicated primary | Haplotigs (no overlap filtering) | Haplotigs (overlaps filtered) |
| --- | --- | --- | --- |
| Total | 77.83 | 85.65 | 85.60 |
| LTR | 75.07 | 83.01 | 82.95 |
| DNA | 0.26 | 0.26 | 0.26 |
| LINE | 0.06 | 0.05 | 0.05 |
| SINE | 0.00 | 0.00 | 0.00 |
| Simple | 2.19 | 2.08 | 2.09 |
| Mobile element | 0.25 | 0.25 | 0.25 |
| rRNA | 0.01 | 0.01 | 0.01 |
| Other | 0.03 | 0.04 | 0.04 |
| Non-repeat | 22.17 | 14.35 | 14.40 |

We estimated a linkage map using dwarf mapping population [52] containing 2,871 SNP markers across 10 linkage groups (Figure 3a). Using the linkage groups, we mapped associate contigs, heterozygosity, SNP, gene, and LTR density to ordered primary contigs to visualize large-scale structural patterns and variation (Figure 3b). Despite performing gene prediction on the masked assembly, we observed that gene and LTR density are moderately correlated on a per-Mb scale (Pearson correlation 0.60, p-value 0.0) (Supplementary Figure 2). To visualize structural variation between primary and associate contigs, we used Gbrowse_syn (Figure 3c).

**Figure 3.**
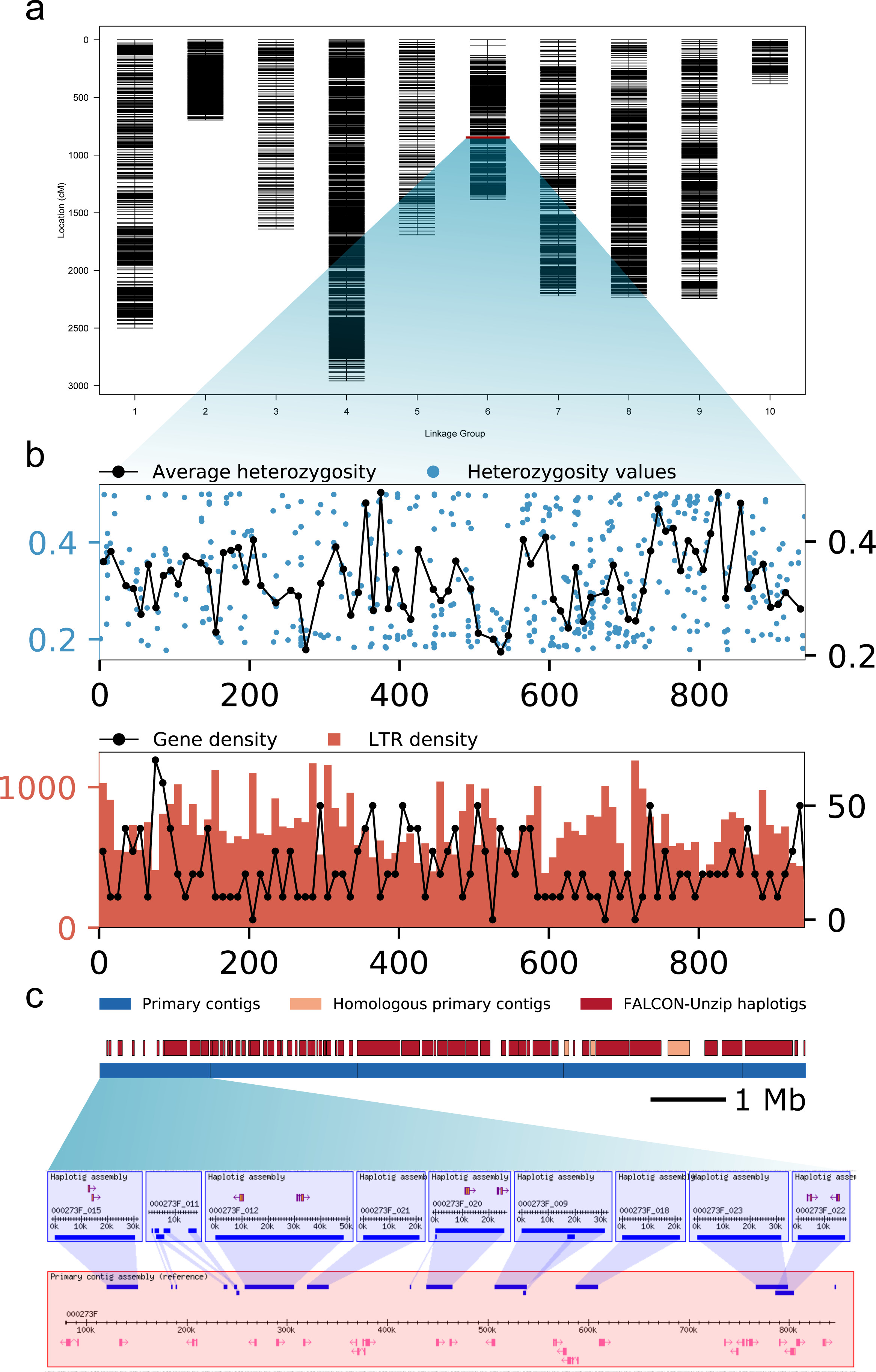

The masked, deduplicated assembly was used as a reference for gene prediction with Augustus [53], which initially generated 153,703 gene models in the primary assembly and 65,274 gene models in the haplotig assembly. After filtering for the longest transcript per gene, requiring protein transcripts to be longer than 150 amino acids [54], and filtering gene models based on homology to repeat-associated Pfam domains, 82,090 gene models remained in the primary assembly and 31,304 gene models remained in the haplotig assembly (Table 4). Repeat-filtering was performed by identifying homology to a set of 91 transposon-associated Pfam domains. There were 9,168 genes featuring homology only to a repeat-associated domain and 5,843 genes featuring homology to both repeat- and non-repeat-associated domains (Table 4). Genes featuring homology only to a repeat-associated domain were removed from further analysis.

**Table 4:** Gene prediction and annotation

| Number of genes | Deduplicated primary | Associate (haplotigs + HPCs) |
| --- | --- | --- |
| Total | 153703 | 65274 |
| Genes encoding proteins with length above 150aa | 91258 | 34625 |
| Genes with a repeat-associated domain | 9168 | 3321 |
| Genes with repeat and non-repeat-associated domains (not removed from assembly) | 5843 | 2344 |
| After filtering repeat-associated genes | 82090 | 31304 |
| Significant hit to UniProt (no repeat filtering applied) | 41092 | 15860 |
| Significant hit to UniProt (repeat filtering applied) | 38324 | 15075 |
| Non-redundant hit to UniProt (no repeat filtering applied) | 10338 | 6447 |
| Non-redundant hit to UniProt (repeat filtering applied) | 10321 | 6429 |

Of the nearly 19,000 repeat-filtered primary genes that overlap haplotigs, 8,085 feature a significant hit to a UniProt gene, but only 4,121 of those significant hits are unique. There are 7,649 genes overlapping HPC regions, and while 3,278 of those genes have a significant hit to UniProt, only 2,038 of the UniProt genes are unique, suggesting widespread duplication of gene function in both cases (Supplementary Table 4).

One of the genes containing both repeat- and non-repeat-associated domains was a putative cannabidiolic acid synthase (CBDAS) gene, previously found only in C. *sativa.* Hop and C. *sativa* share localized regions of conserved synteny of genes involved in cannabinoid synthesis (Figure 4a). Multiple putative BBE-like, CBDAS, and CBDAS-like hop genes share conserved synteny with genes on chromosome nine of C. *sativa* (Figure 4a,b), which contains tandemly repeated CBDAS genes nested within LTRs [32]. We also identified a region of co-localized genes featuring berberine bridge enzyme (BBE)-like genes (Figure 4b). The putative CBDAS gene contains three domains of unknown function (DUF), a transposase domain, and a FAD-binding domain (Figure 4c). Only the region of the gene overlapping the FAD-binding domain, corresponding to where CBDAS homology occurs, shows evidence of expression in our RNA-seq data (Figure 4a). Upstream of the expressed region of the gene, an intron contains segments of Gypsy-type LTRs. Further upstream where the transposon domain occurs, however, no repeats are annotated. Although gene prediction was performed on the masked assembly, repetitive regions could have been incompletely masked, which would explain why prediction extended through a repeat sequence. The fusion of a transposon domain and putative CBDAS gene in hop would be a telling consequence of the highly repetitive composition of the genome, and is consistent with a previous finding that CBDAS genes are located near LTRs in C. *sativa* [32, 33]. In total, we found colocalized regions of genes within three linkage groups in hop and three chromosomes of C. *sativa*.

**Figure 4.**
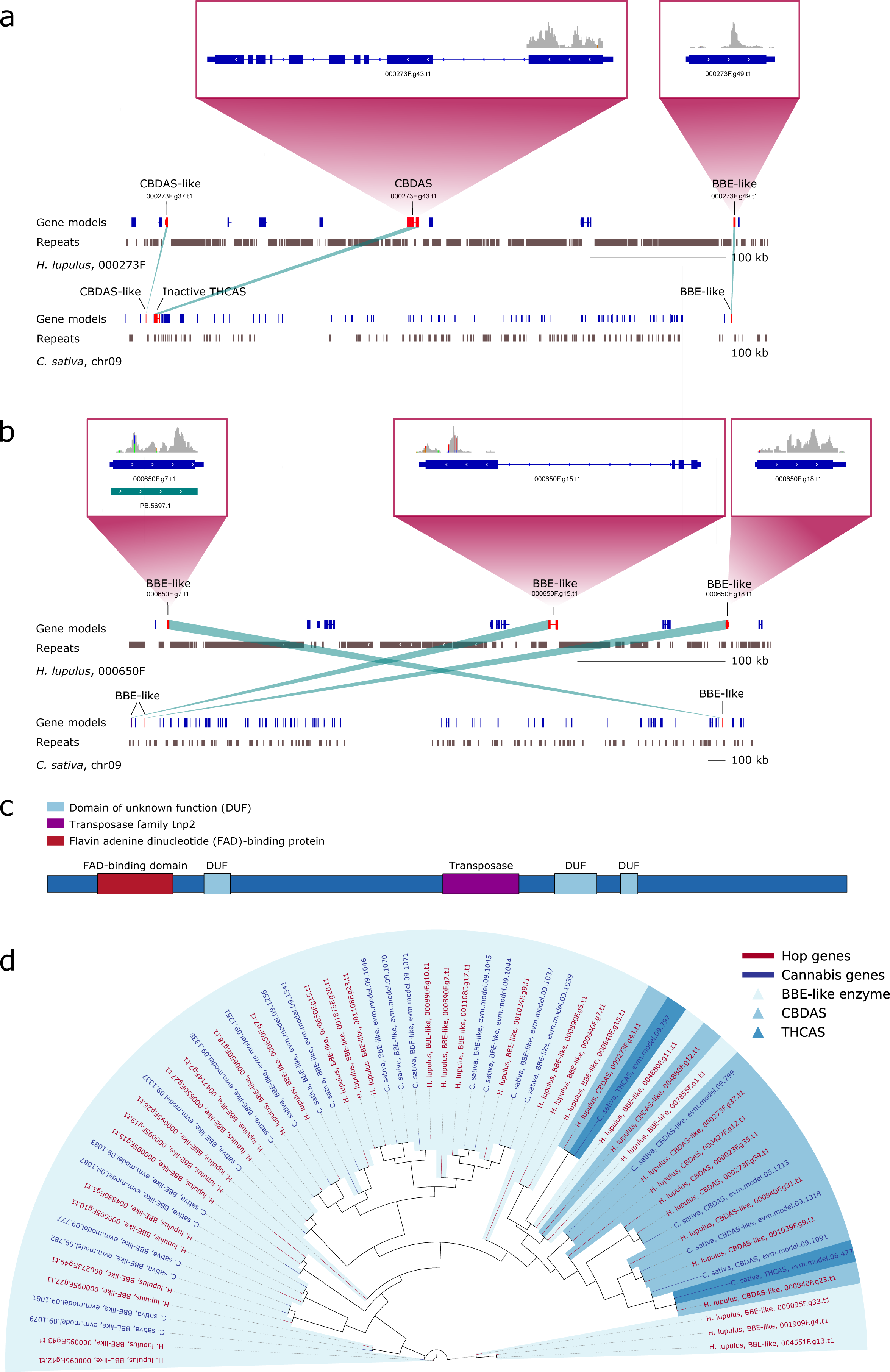

To identify hop and C. *sativa* genes homologous to CBDAS, tetrahydrocannabinolic acid synthase (THCAS and THCAS-like), and BBE-like genes, we first aligned all hop and C. *sativa* peptides to a set of UniProt Embryophyta sequences. We collected peptide sequences with highest scoring blastp alignments to either CBDAS, THCAS, or BBE-like genes. A phylogenetic tree was constructed by aligning these genes from hop and C. *sativa* (Figure 4d). Synteny was assessed by performing a mutual best hit (MBH) analysis to identify genes in hop and C. *sativa* with highest shared homology to CBDAS, CBDAS-like, THCAS, THCAS-like, and BBE-like genes. Additionally, to investigate the extent of gene duplication in hop and C. *sativa,* we computed the number of occurrences of significant homology to Uniprot Embryophyta genes. We found that while C. *sativa* is enriched for single-copy genes, hop shows enrichment relative to C. *sativa* for two and four gene copies (Supplementary Figure 3), providing supporting evidence for WGD in hop.

Large-scale structural variation between primary contigs and their associate contigs was visualized with mummerplot (Figure 5a, Supplementary Figure 4a). Three out of eight of the associate contigs aligning to primary contig 000040F are HPCs (Figure 5a), which exhibit structural variation, including inversions (HPCs 003851F and 003705F). Primary contig 000145F features 14 associate contigs, and all but one are haplotigs. The single HPC (004045F) covers the middle of the primary contig, and features more variation and shorter alignment blocks than the haplotigs (Supplementary Figure 4a). We observe that HPCs have greater structural variation than haplotigs, which are generally shorter than HPCs and have near-perfect alignments to their primary contig. Overall, alignment block length and coverage is longer in haplotigs than in HPCs (Supplementary Figure 4b,c).

SNPs, insertions, deletions, and structural variation are more abundant in HPCs than haplotigs (Figure 5b), and gap lengths are longer and more abundant in primary contig and HPC alignment blocks (Figure 5c). We used Kimura 80 distance (55], an evolutionary distance that is a function of the rate of transitions and transversions, to assess sequence divergence between haplotypes. This distance is larger between primary contigs and HPCs than primary contigs and haplotigs (Figure 5d). The conditional probability associated with transitions between haplotypes is higher in the alignment blocks of primary contigs and HPCs than primary contigs and haplotigs (Figure 5e). We also found a weak correlation between the heterozygosity evaluated from 660 individuals and Kimura distance between haplotypes, as expected (Supplementary Figure 5).

**Figure 5.**
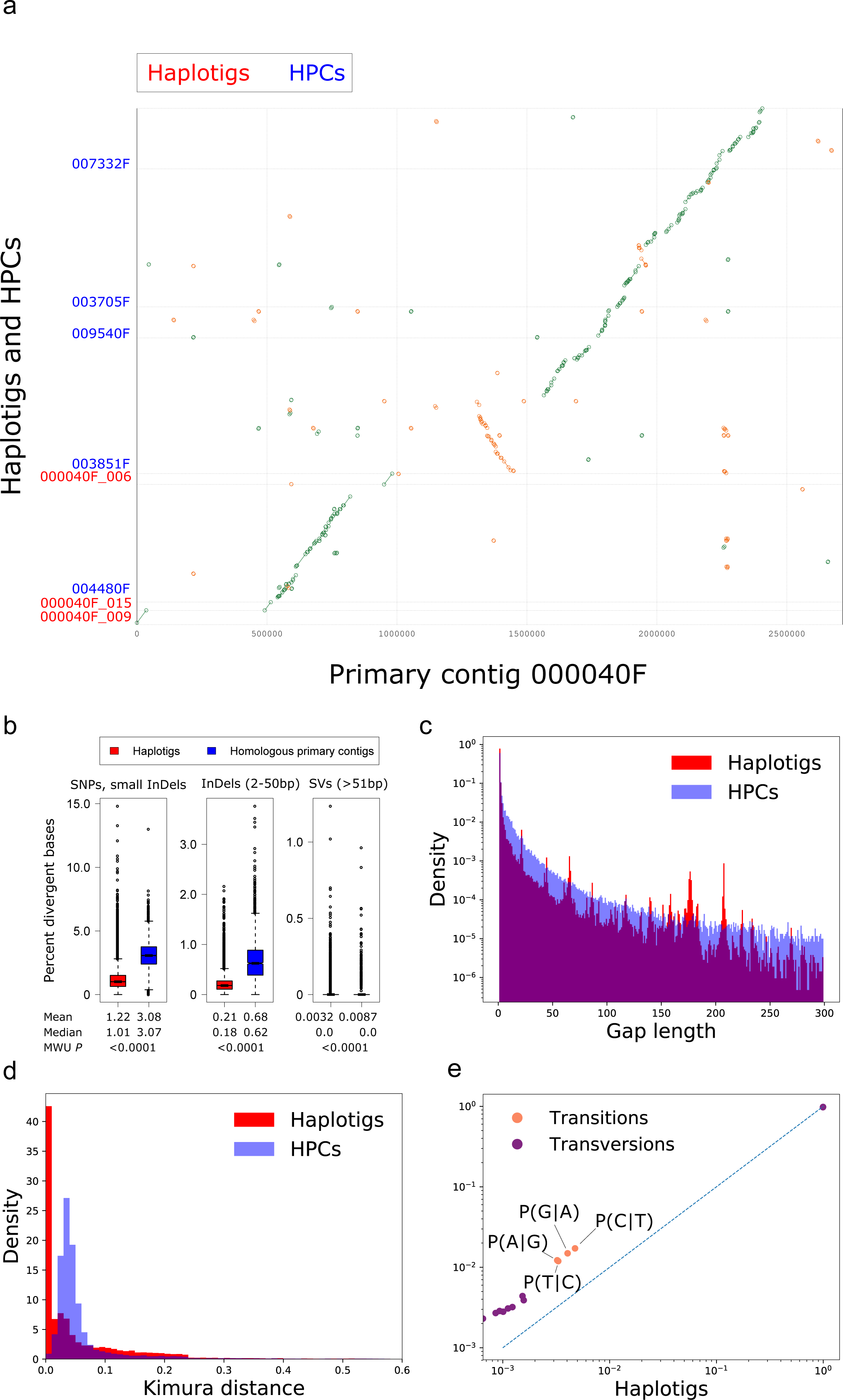

Having found no significant difference in the LTR content of HPCs and haplotigs, we further investigated divergence due to gene content. To assess divergence between the genes of primary and associate contigs, we developed an approach for extracting projected gene sequences from the primary assembly to the associate contig assembly. We first aligned the primary contigs and associate contigs, and then extracted and concatenated exon sequences from both primary contig and associate contig alignment blocks based on the coordinates of the gene model in the primary contig. The length difference between coding sequences (CDS) of primary and associate contigs was computed, revealing a core set of ∼30,000 genes without any gaps. Although ∼80,000 genes were predicted in the primary assembly, the presence of only ∼30,000 genes containing no gaps in alignment suggests that a core set of genes are strongly conserved between primary contigs and haplotigs (Figure 6a). In addition, we found an enrichment of CDS sequences with gaps totaling a multiple of three, indicating a conserved frame (Figure 6a).

**Figure 6.**
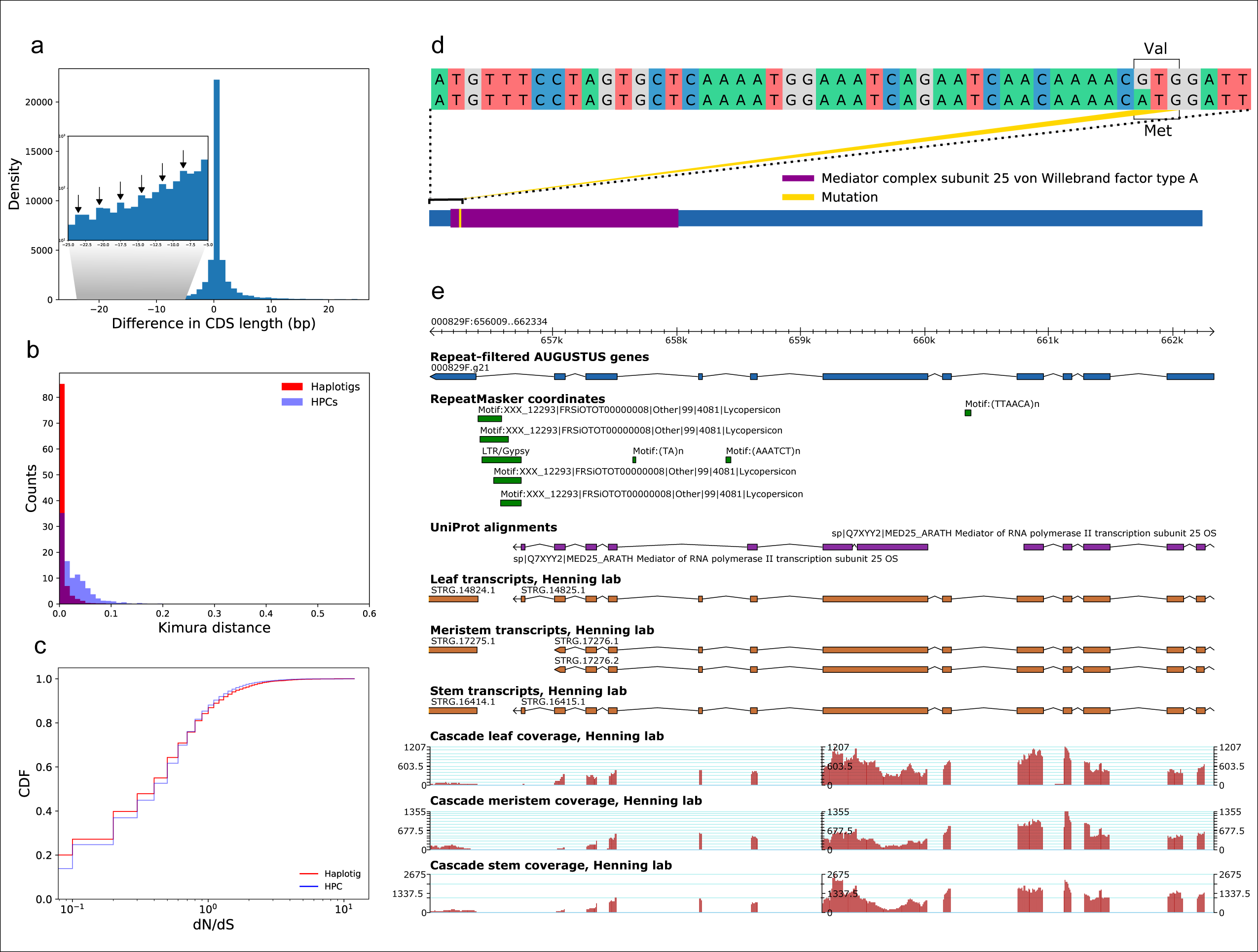

We then investigated functional enrichment (Table 5) and mutation rates of genes in haplotigs and HPCs as a mechanism for divergence, under the hypothesis that cultivation and environmental stresses have resulted in greater divergence of certain gene regions. By comparing CDS regions between primary and associate contigs, we found that HPCs and primary contigs have a greater Kimura 80 distance than primary contigs and haplotigs (Figure 6b). To further assess the biological significance of the greater divergence in HPCs compared to haplotigs, we analyzed GO term enrichment in the most diverged genes within HPCs. We collected genes within HPC alignments that are equally or more diverged than the top 25% of haplotig alignment genes, based on Kimura distances. We performed a binomial test to ascertain whether the number of genes associated with GO term enrichment in HPCs is significantly greater than the expected rate of occurrence computed from genes above the 25% Kimura distance threshold within haplotigs. We calculated the probability p of GO term occurrence in haplotigs, the observed occurrence of a given GO term *k* in the HPCs, and the total number of HPC genes n with an associated GO term above the Kimura distance threshold (Supplementary Figure 6). An expected probability for HPCs was calculated by multiplying n and *p.* A binomial test was performed on only those GO terms with an observed *k* above the expected probability. Enriched biological functions include 27 genes associated with carbohydrate metabolism (GO:0005975: carbohydrate metabolic process), seven genes involved in chitin signaling (GO:0010200: response to chitin), and several genes related to biotic stress (GO:0009615: response to virus) (Supplementary Table 5).

**Table 5:** Enriched GO Terms in haplotigs and HPCs

| GO term | GO description | FDR | Observed | Expected |
| --- | --- | --- | --- | --- |
| Biological Processes GO terms enriched in haplotigs |  |  |  |  |
| GO:0010038 | response to metal ion | 0.002 | 20.000 | 6.890 |
| GO:0060320 | rejection of self pollen | 0.002 | 16.000 | 5.100 |
| GO:0009627 | systemic acquired resistance | 0.022 | 22.000 | 10.200 |
| GO:0006506 | GPI anchor biosynthetic process | 0.022 | 6.000 | 1.280 |
| GO:0009737 | response to abscisic acid | 0.046 | 179.000 | 142.000 |
| Biological Processes GO terms enriched in HPCs |  |  |  |  |
| GO:0006397 | mRNA processing | 0.024 | 62.000 | 38.000 |
| GO:0010152 | pollen maturation | 0.032 | 7.000 | 1.900 |
| GO:0009685 | gibberellin metabolic process | 0.032 | 3.000 | 0.399 |
| GO:0001578 | microtubule bundle formation | 0.032 | 2.000 | 0.169 |
| GO:0009609 | response to symbiotic bacterium | 0.032 | 2.000 | 0.153 |
| GO:0007049 | cell cycle | 0.039 | 36.000 | 21.200 |
| Cellular Components GO terms enriched in haplotigs |  |  |  |  |
| GO:0000111 | nucleotide-excision repair factor 2 complex | 0.000 | 3.000 | 0.113 |
| Cellular Components GO terms enriched in HPCs |  |  |  |  |
| GO:0005680 | anaphase-promoting complex | 0.032 | 4.000 | 0.583 |
| GO:0009509 | chromoplast | 0.045 | 3.000 | 0.505 |
| GO:0000775 | chromosome, centromeric region | 0.045 | 6.000 | 1.640 |
| GO:0005652 | nuclear lamina | 0.045 | 2.000 | 0.205 |
| Molecular Function GO terms enriched in haplotigs |  |  |  |  |
| None |  |  |  |  |
| Molecular Function GO terms enriched in HPCs |  |  |  |  |
| GO:0004867 | serine-type endopeptidase inhibitor activity | 0.009 | 7.000 | 1.250 |
| GO:0005324 | long-chain fatty acid transporter activity | 0.010 | 3.000 | 0.240 |
| GO:0004866 | endopeptidase inhibitor activity | 0.011 | 2.000 | 0.128 |
| GO:0004108 | citrate (Si)-synthase activity | 0.011 | 3.000 | 0.305 |
| GO:0005516 | calmodulin binding | 0.011 | 75.000 | 49.300 |
| GO:0004560 | alpha-L-fucosidase activity | 0.012 | 4.000 | 0.593 |
| GO:0004555 | alpha, alpha-trehalase activity | 0.016 | 2.000 | 0.160 |
| GO:0003876 | AMP deaminase activity | 0.022 | 2.000 | 0.192 |
| GO:0003777 | microtubule motor activity | 0.022 | 23.000 | 11.500 |
| GO:0000146 | microfilament motor activity | 0.046 | 3.000 | 0.529 |

Investigating further into the types of mutations, we examined the rate ratio of non-synonymous mutations relative to synonymous mutations (dN/dS). HPCs contain more genes with dN/dS >1 relative to haplotigs (Figure 6c), indicating a subset of genes in the HPCs under greater positive selection. To evaluate whether the genes under positive selection are enriched for specific gene functions, we assessed GO term enrichment with a hypergeometric test, followed by a Benjamini-Hochberg multiple test correction (FDR:50.05). We compared the set of hop genes featuring dN/dS >1 to the set of all genes with an associated GO term. Genes with dN/dS >1 are enriched for Biological Processes GO terms associated with stress response (GO:0009693: ethylene biosynthetic process, GO:0009267: cellular response to starvation, GO:0009411: response to UV, and GO:0010569: regulation of double-strand break repair via homologous recombination) (Supplementary Table 6). Other genes under positive selection are involved in growth and development (GO:0010091: trichome branching and GO:0009800: cinnamic acid biosynthetic process). Based on GO term enrichment, genes with dN/dS > 1 are under greater selective pressure in the presence of abiotic and biotic stresses. We investigated one of the genes with dN/dS > 1 (000829F.g21.t2:656009-662333) featuring an enriched Biological Processes GO term, “trichome branching.” This gene features a mutation in its protein domain, “Mediator complex subunit 25 von Willebrand factor type A” (Figure 6d), and provides an example of LTR sequences located within an intron (Figure 6e).

## Discussion

The large size and high heterozygosity of the hop genome has challenged previous efforts at assembly. As a result, many questions regarding the hop genome have remained unanswered. In this study, we have used a haplotype-phased assembly to overcome these challenges, and to investigate repeat regions, gene duplication, and structural variation between haplotypes. We have presented a deduplication strategy to effectively identify and reassign HPCs, resulting in a refined, phased diploid assembly that contains a comprehensive repertoire of genes and captures other relevant genomic features. The haplotype-phased assembly offers a unique opportunity to study structural variation between the haplotype assemblies, and provides insight into the factors influencing the evolution and selection of hop.

Short-read assembly fragmentation has presented obstacles in estimating the repeat content of the hop genome. The Shinsuwase assembly is 34.7% repetitive with LTRs comprising 32.3% of total scaffold sequence [5]. More recently, genome repeat content was estimated at ∼60% using genome shotgun sequencing reads from hop, as well as Japanese wild hop, var. Lupulus and var. cordifolius [42]. Our long-read assembly has allowed for a more-accurate estimation of repeat content, suggesting that hop is 78% repetitive, comparable to maize (Figure 2a,b). This finding has implications for understanding the evolution of the Cannabaceae family (Figure 1a).

Hop and C. *sativa* diverged ∼21 million years ago [56]. Hop is four times larger than C. *sativa,* and the significant difference in genome size is hypothesized to be a result of WGD [57]. The genome of *Humulus Jupulus* is twice as large as *Humulus japonicas,* although minimal information is known about *H. japonicus.* Our finding suggests that hop and C. *sativa* contain a similar percentage of repeat content, as C. *sativa* is ∼73% repetitive [42]. The similar repeat percentage between hop and C. *sativa* supports the hypothesis that hop genome expansion is a consequence of WGD followed by a return to a diploid state through rearrangement and gene loss [57, 58], rather than LTR-driven expansion observed in other plants [59]. Hop is enriched for two and four gene copies relative to C. *sativa* (Supplementary Figure 3), supporting the WGD hypothesis.

Hop and C. *sativa* share common enzymes, including polyketide synthases and prenyltransferases, that participate in the synthesis of key compounds [60], and our much improved assembly will help to elucidate the evolution of these enzymes. To our knowledge, we are the first to report the presence of CDBAS in hop, which may have previously been obscured due to proximity to LTRs. CBDAS is the hypothesized common ancestor of CBDAS and THCAS [61] due to high sequence similarity (84%) [62]. The presence of CBDAS in hop supports the hypothesis that CBDAS is the ancestral gene of CBDAS and THCAS, because CBDAS would be present in the common ancestor. The emergence of THCAS is thought to be a result of CBDAS duplication and diversification [61], possibly copied and dispersed by LTRs [32]. Both synthases also share high sequence similarity with BBE-like enzymes, which catalyze oxidation reactions and are highly abundant in plants [63].

One of the hop genes containing both repeat- and non-repeat-associated domains has significant homology to CBDAS (Figure 4a). Among the ten predicted exons of the putative CBDAS gene in hop (000273F.g43.t1-region 1034766 - 1043385) only the exon with strong homology to CBDAS is expressed; therefore, the fusion could be a gene-prediction artifact. The putative CBDAS hop gene is expressed in lupulin gland, leaf, and meristem tissues, with highest expression occurring in leaf. The presence of a putative CBDAS gene near LTR sequences in hop provides further justification for a more-complete assembly of the genome, especially since an abundance of CBDAS homologs within LTR regions in C. *sativa* was previously recognized. Further work will be necessary to refine the gene models containing both repeat and non-repeat-associated domains, and to determine whether hop produces detectable amounts of CBD at any point during flower development.

The large number of primary assembly gene models (82,090 after repeat-filtering) could be a result of residual primary contig duplication. However, recent assemblies of barley (∼5.3Gb), *Aegilops tauschii* (4.3Gb), and *Nicotiana tabacum* (4.5Gb) each contained >80,000 gene models prior to filtering for high-confidence genes [64–66] indicating that a very large number of genes predicted from large and complex genomes is not inconceivable. Of the 82,090 gene models in the primary assembly, 38,324 feature a significant hit to a UniProt gene and 10,321 of those hits to UniProt are non-redundant, indicating that significant gene duplication is widespread (Table 4). From the projected gene models extracted from alignment blocks, ∼30,000 contain no gaps in alignment, suggesting a core set of genes are more conserved between primary and associate contigs (Figure 6a). The Shinsuwase cultivar assembly contains 22,201 annotated protein-coding genes [5] and the Teamaker cultivar assembly contains 16,161 annotated protein-coding genes [6] after filtering for repeat-associated protein-coding genes. Future work will need to integrate these data and produce a refined list of hop genes.

The haplotypes are mostly well conserved, but are punctuated by regions of high heterozygosity that are enriched for genes under positive selection. Genes in HPCs are characterized by a larger evolutionary distance and greater variation compared to haplotigs, and are enriched for genes involved in growth, development, and response to abiotic and biotic stresses. The abundance of genes associated with these processes, as identified by Kimura distance and dN/dS analysis, suggests a genomic landscape reacting to shifting abiotic and biotic stress conditions. This finding, combined with the heavily duplicated gene composition of the genome, illustrates the response of the genome to environmental stresses through time. Investigation of haplotype variation provides insight into the patterns of selection that separate the English-Russian maternal pedigree from the USDA paternal pedigree of Cascade.

Our successfully deduplicated assembly sets a foundation for chromosome-level scaffolding with a high-throughput chromatin conformation capture (Hi-C) library, which will further refine the assembly by providing long-range DNA contact information. An expanded assembly capturing regions beyond gene space will also enable future work on enhancers and regulatory DNA that were obscured due to fragmentation. Going forward, a chromosome-level assembly will provide a foundation for investigating the occurrence of WGD in hop [42], and to understand how duplication, repeat-expansion, and conserved synteny have driven biosynthesis and disease resistance in hop.

## Methods

### Sample Collection and DNA Sequencing

To prepare for sequencing, multiple samples of approximately 100 µg of young leaves from Cascade were collected and placed on ice, and DNA was extracted using a Qiagen DNeasy miniprep kit with some modifications. To prevent shearing, chemical precipitation and glass hooks were used instead of spin columns. Two SMRTbell libraries were constructed from DNA with a required minimum length of 10kb. Sequencing was performed on PacBioRS II with P6-C4 chemistry and Sequel 1.2.1-2.0 chemistry. For the draft assembly, total raw read size was 288 Gb, 182 Gb for seed reads, and 135 Gb for the pre-assembled reads (p-reads), with low-coverage regions removed during assembly. Only subreads greater than 500bp and p-reads longer than 11kb were used.

### Genome Assembly and Phasing

The goal of a phased, diploid assembly is to produce an assembly for each haplotype of the diploid genome. Genome assembly and phasing proceeded in five stages: pre-assembly, overlap, FALCON, FALCON-Unzip, and phased polish. Long-read sequences generated by SMRT sequencing were assembled with FALCON, a diploid-aware, *de novo* assembler [46]. The haplotypes were phased by FALCON-Unzip based on structural variants (SV) and single-nucleotide polymorphisms (SNPs) in the genome. The output of FALCON-Unzip included a set of primary and associate contigs, corresponding to an assembly of both haplotypes of the diploid genome.

Error-correction was performed during pre-assembly. Reads longer than 10kb were selected as seed reads, and all shorter reads were aligned to the seed reads to generate p-reads, which are high-accuracy consensus sequences. Pre-assembly yield was 75%, and served as a metric of data quality and coverage. Next, p-reads were aligned to each other and assembled into contigs. Polishing consisted of aligning all subreads to the draft contigs to generate final contig consensus sequences.

In areas of low heterozygosity, haplotypes are collapsed into a single primary contig. Areas of medium heterozygosity, with less than 4% structural variation, are used for phasing and identification of haplotigs. Highly heterozygous regions that are too diverged to be identified as homologous are assembled as independent primary contigs [46], resulting in duplicated primary contigs. As independently-assembled homologous primary contigs (HPCs) are too diverged to be assembled as haplotypes, further steps must be taken to identify and reassign HPCs as associate contigs.

### Deduplication with purge_haplotigs

HPCs were identified by read coverage and sequence alignment with purge_haplotigs [67L._Duplicated primary contigs were expected to have half read-depth because reads would be split between multiple contigs. Candidate HPCs featuring half read-depth were analyzed for homology with blastn [68] and LASTZ [69]. Alignment statistics were computed to differentiate between highly repetitive alignments and homologous regions.

### Assessment of Deduplication

The deduplication strategy was assessed by Benchmarking Universal Single-Copy Orthologs (BUSCO) [70], which consists of a highly-conserved set of genes that are expected to be present in single-copy across closely-related organisms. BUSCO provides a measure of the extent of duplication and completeness of gene content in an assembly. The purge_haplotigs-deduplicated assembly contained 451 duplicated BUSCO genes, suggesting that further deduplication was possible. The goal moving forward was to use BUSCO to identify which alignment and filtering parameters maximized the number of single-copy genes and minimized the number of duplicated, fragmented, and missing genes.

### Further Sequence Alignment-based Deduplication

Following deduplication with purge_haplotigs, the assembly was still larger than expected and a large number of duplicated BUSCO genes persisted. We further developed a sequence alignment-based strategy to identify HPC pairs that resisted identification by purge_haplotigs.

The first step was to identify homologous pairs using a fast, computationally inexpensive method. Homologous regions between contigs were retrieved from megablast alignments between all primary contigs. Megablast output was sorted by score in descending order, discarding self-hits. Top, unique hits were stored, imposing an E-value of less than 1e-100, while redundant, significantly overlapping hits were discarded. A position-specific hit count along the length of the primary contig was computed, ultimately providing an average hit count for the entire length of the contig. Contigs aligning to more than 10 contigs on average were removed. The hit count corresponded to a contig pair having high homology and low occurrence of duplication. Contig pairs with a hit count above the threshold were selected for further analysis with pairwise DNA alignment tools, LASTZ and MUMmer. Visualization of MUMmer alignments with Mummerplot allowed for verification of alignment strength, as well as identification of large-scale structural features such as tandem repeats and inversions.

To assess the strength of homology between contigs, the alignment density between primary contig pairs was computed. Density values were calculated by summing the alignment score or coverage to get a total score or coverage, and then dividing by the length of the shorter contig. The threshold density value for filtering was selected based on the correspondence of score or coverage density to homology between primary contigs, as validated by visual inspection with Mummerplot. Filtering parameters included minimum coverage, score density, continuity, and identity set to 20%. Maximum percent overhang was a filtering parameter where the maximum percentage of the shorter contig that remained unaligned at its beginning or end to the longer contig was set to 40%. Primary contigs featuring more than 40% maximum overhang were removed from further deduplication analysis.

Some contig pairs exhibited a misleadingly large score sum and score density in the presence of redundant, overlapping alignment blocks. By requiring that alignment blocks not overlap by more than 50% relative to the shorter contig, a score sum based on fewer redundant and overlapping alignment blocks was calculated.

Identification of strong homology was often confounded in the presence of short, spurious, off-diagonal alignment blocks, with the result being erroneously inflated coverage density values. Alignment blocks from Mummer were clustered based on their proximity to remove short and distant alignment blocks. Alignment blocks were grouped into clusters if the block start positions occurred within 10 kb of each other. If the start positions were separated by more than 10 kb, a new cluster was initialized. Clusters overlapping by less than 50% were stored. For clusters overlapping by more than 50%, the larger of the clusters was retained. Also, only clusters containing five or more alignment blocks were stored to reduce the number of short, spurious, off-diagonal hits, thereby preventing erroneously high-coverage densities. Minimum cluster length was set to 20kb.

To achieve the total coverage density of all clusters combined, the start position of the first alignment block in the cluster was subtracted from the end position of the last alignment block in the cluster, and the difference was divided by the length of the shorter contig. A cluster coverage density threshold above 25% was imposed, signifying that the total span of cluster coverage should cover at least 25% of the shorter primary contig. All values were calculated relative to the shorter contig.

Once high-confidence HPCs were identified, they were reassigned to the associate contig assembly along with the haplotigs. Both purge_haplotigs [67] and our sequence alignment-based deduplication strategy captured a slightly different set of HPCs. HPCs identified by purge_haplotigs were given precedence above the sequence alignment-identified HPCs. HPCs identified by the sequence alignment-based pipeline were reassigned as associate contigs if they had not been identified by purge_haplotigs as HPCs. By integrating the non-overlapping pairs identified by both pipelines, further recovery of associate contigs was achieved.

Many FALCON-Unzip haplotigs and HPCs identified by purge_haplotigs overlapped in position on their corresponding primary contig. The shorter of the overlapping FALCON-Unzip haplotigs and HPCs were removed from the haplotig assembly to prevent redundancy in calculating statistics.

### Repeat prediction and annotation

The primary contig and haplotig assemblies were masked with RepeatMasker [51] using a database of Eudicot repeat elements from MIPS PlantDB [SO]. *De nova* identification of LTRs was done with LTR_FINDER [47], LTRharvest [48], and LTR_retriever [49]. The resulting combined, masked assembly was used for gene prediction.

### RNA-seq preparation and transcriptome assembly

RNA-seq libraries from hop tissues were generated using RNeasy Plant Mini Kit (Qiagen), using glass hooks to remove RNA from tubes. Sequencing was performed on an Illumina HiSeq 3000 with 100 bp stranded libraries. Three TruSeq RNA (Qiagen) libraries were developed from leaf, stem, and meristem tissues with each library sequenced on a single lane. The transcriptome assembly was generated by aligning RNA-seq to the deduplicated primary contig assembly with Hisat2 [71], followed by assembly into transcripts with Stringtie [72]. Protein-coding sequences were predicted with Transdecoder [73] and only the longest open reading frame (ORF) sequences were used for further analysis.

### Gene prediction

Multiple sources of evidence were incorporated as hints to Augustus for gene prediction [74]. Hints were derived from ESTs, Embryophyta genes from UniProt, predicted protein-coding transcripts from hop, and hop RNA-seq. Hop ESTs from TrichOME [75] and NCBI were aligned to the masked, deduplicated assembly using blastn. Next, all ESTs with a hit to the assembly were aligned to the assembly with exonerate [76]. UniProt genes were aligned to the masked, deduplicated assembly with blastx, and then all UniProt genes with a hit to the assembly were aligned to the assembly with exonerate. Protein-coding transcripts generated by Transdecoder were aligned to the masked, deduplicated assembly with exonerate.

### Inparanoid clustering analysis

Peptide sequences coded by canonical transcripts from 115 species covering a range of clades from the tree of life (Supplementary Table 2) were downloaded from respective sources. The sequences were provided to the gene orthology prediction workflow involving Inparanoid [77] as previously described [78]. Gene family clusters were queried further to find unique and overlapping genes between the five gene sets from *A. thaliana, V. vinifera,* C. *sativa,* as well as the primary and haplotig gene models. The unique gene clusters identified in the two hop assemblies were queried for potential GO [79], biological pathways, and InterProScan domain [80] association.

### Analysis of coding sequences

Haplotigs were aligned to all primary contigs using LASTZ. Gene model sequences were extracted from primary contig alignment blocks based on genomic coordinates of the gene models, allowing the presence of gaps. Gene model coordinates in the primary contigs were projected to the corresponding positions in the haplotig alignment blocks, using the genome alignment as a guide. CDS from primary contig and haplotig alignment blocks were then extracted. CDS alignments were processed with MACSE [81], using the exportAlignment option. Internal stop codons were denoted as ‘NNN’ and codons containing frameshifts were denoted as ‘---.’ Further processing was performed to remove frameshift-containing and stop codons.

### Variation between haplotypes: dN/dS, gap lengths and rates, Kimura distance, and GO term enrichment

Variation between haplotypes was assessed by calculating mutation and gap rates, gap lengths, Kimura 80 distance, and the non-synonymous to synonymous substitution rate ratio (dN/dS).

To investigate functional enrichment of HPC genes, we first identified haplotig genes with the top 25% of Kimura distances. The lowest value in the top 25% of Kimura distances was used to set the distance threshold for the HPC genes. We performed a binomial test to assess whether GO term enrichment of genes in HPCs with a Kimura distance above the distance threshold is significantly greater than the expected rate of GO term occurrence among haplotig genes with the top 25% of Kimura distances. Probability p was calculated from haplotigs, while k was the observed occurrence of a given GO term in the HPCs, and n was the total number of HPC genes with an associated GO term above the Kimura distance threshold.

Calculation of dN/dS provides a method to quantify selection within protein-coding regions, treating the two haplotypes as separate individuals. The dN/dS ratio was computed between pairs of CDS from the primary contigs and corresponding associate contigs. Genes from the primary assembly were aligned to a set of 37,364 Embryophyta genes from UniProt with blastp (E-value::51e-5) and MBHs were identified. Functional enrichment of genes under positive selection was assessed by comparing the set of genes with dN/dS > 1 to all genes with an associated GO term. To test for enrichment, a hypergeometric test was performed, followed by a Benjamini-Hochberg multiple test correction (FDR::50.05). Gap rates and lengths, as well as Kimura 80 distance, were calculated from the alignment blocks between primary contigs and haplotigs.

### Phylogenetic tree and synteny analysis

To assess homology and evolutionary distance between hop and C. *sativa,* a phylogenetic tree was constructed. All hop protein sequences were aligned to the set of 37,364 Embryophyta genes from UniProt with blastp (E-value::51e-5), and the same step was taken with the protein sequences from C. *sativa.* The extent of gene duplication among hop and C. *sativa* genes was assessed by counting the number of times a UniProt gene featured significant homology to a hop or C. *sativa* gene. An enrichment value was computed by subtracting the number of C. *sativa* hits from the number of hop hits, and then dividing by the number of C. *sativa* hits plus one. The enrichment value provided a measure of the relative abundance of gene copy number variation between hop and C. *sativa*.

Significant hits to cannabidiolic acid synthase (CBDAS and CBDAS-like) genes, berberine bridge enzyme-like genes (BBE-like), and tetrahydrocannabinolic acid synthase (THCAS and THCAS-like) genes were extracted for both hop and C. *sativa,* and these hits were aligned to each other. A phylogenetic tree was constructed from the top hits between hop and C. *sativa* genes. The phylogenetic tree was constructed using ClustalW [82], trimAL [83], PhyML [84], and the python module ETE3 [85].

MBHs were identified from the set of hop and C. *sativa* protein sequences with a significant hit to CBDAS, CBDAS-like, THCA, THCA-like, and BBE-like genes. Several syntenic regions featuring genes from chromosome nine of the high-CBDA cultivar were investigated. Synteny was visualized with Integrative Genomics Viewer (IGV) [86].

### SNP identification

Plant material, DNA extraction, library prep, sequencing, and identification of SNP markers for developing a linkage map were previously reported [52] except for using the haplotype-phased assembly as reference genome. Over one million SNPs were identified across 660 unique genotypes, including both cultivars and USDA experimental varieties. Genotype-by-sequencing (GBS) data from 94 offspring and parents for a population study to identify alleles associated with short-stature (dwarf mapping) was extracted from this global data set and further filtered down to markers with 2X coverage and presence in 95% of all mapping population and parents. GBS data from this data set were exported in hapmap format for linkage map development.

### Linkage map

Estimation of a linkage map for a dwarf mapping population followed the same process as previously reported [52]. The starting number of highly filtered markers was approximately 13,000. We developed linkage maps for both parents and then integrated both maps into a single consensus map using the same methods as previously reported. The final map contained 2,871 SNP markers across 10 linkage groups. As we had identified the sex of all members of the mapping population we were able to identify the pseudo-autosomal linkage group by mapping sex as a phenotype.

## Abbreviations

Single molecule real-time (SMRT) sequencing; single nucleotide polymorphism (SNP); structural variant (SV); homologous primary contig (HPC); benchmarking universal single-copy orthologs (BUSCO); transposable element (TE); long terminal retrotransposon (LTR); whole genome duplication (WGD); cannabidiolic acid synthase (CBDAS); berberine bridge enzyme-like (BBE-like); tetrahydrocannabinolic acid synthase (THCAS)

## Declarations

### Availability of data and materials

The datasets generated and analyzed in this study are available at HopBase.org and NCBI under the BioProject ID PRJNA562558. Scripts are available at: https://github.com/padgittl/CascadeHopAssembly/.

### Funding

Funding for sequencing was provided by PacBio and Sierra Nevada Brewing Company. This work was supported by a grant from USDA (USDA-ARS CRIS Project 2072 21000 051 00D). JE and PJ are partially supported for their gene family work by NSF USA award <u>#105:1340112.</u>

### Conflict of Interest Statement

SK, PP, GC, and DR are full-time employees at Pacific Biosciences, a company developing single-molecule sequencing technologies.

## Notes

http://hopbase.org/

