## Supplementary Materials for "A phased, diploid assembly of the Cascade hop *(Humulus lupulus)* genome reveals patterns of selection and haplotype variation"

4 USDA ARS

5 School of Electrical Engineering and Computer Science, Oregon State University

6 Department of Botany and Plant Pathology, Oregon State University

**Assembly deduplication**

The first step towards identifying homology between primary contigs entailed aligning all primary contigs to each other with megablast. An example command is:

megablast -i contig.fasta -d assembly.fasta -o contig_vs_assembly.outfmt8 -e 1e-5 -m 8 -F F -W 28

Contig pairs with significant homology and low duplication frequency were tagged as candidate HPC pairs and further assessed with LASTZ and mummer. Low duplication frequency was assessed with this command:

python printLowDuplicationFrequencyBlastHits.py outfmt8.megablast.txt > lowDuplicationFrequencyBlastHits.txt

An example LASTZ command is:

lastz HPC.fasta primaryContig.fasta --gfextend --hspthresh=20000 --chain --gapped --output=HPC_vs_primaryContig.txt --format=general:score,name1,strand1,size1,zstart1,end1,name2,strand2,size2,zstart2,end2,identity,continuity,coverage --inner=10000 --identity=80 --strand=plus

An example mummer command is:

mummer -maxmatch -b -c -l 100 primaryContig.fa HPC.fa > primaryContig_vs_HPC.mums

To visualize alignments, mummerplot was used, and an example command is:

mummerplot -postscript -p 007178F_vs_011094F 007178F_vs_011094F.mums

Alignment density and coverage values were obtained from this script:

python computeNonOverlappingScoreDensity.py lastzOutputFileList.txt > results.txt

Candidate HPCs for a given primary contig were compiled into clusters with this script:

perl processLASTZ.pl results.txt primaryContigLengths.txt

Deduplication was performed with this command:

python deduplicateContigs.py clusters.txt primaryContigLengths.txt lastzOutputFileList.txt purge_haplotigs_lastzOutputFileList.txt listOfPrimaryContigs.txt listOfHaplotigs.txt curated.contig_associations.log outBaseName

**Repeat Masking**

*De novo* identification of LTR sequences in hop was performed by first creating a suffix array index for the genome assembly with this command:

gt suffixerator –db primaryAssembly.fasta –indexname hopsindex -tis -suf -lcp -des -ssp –dna

Next, analysis with LTRharvest was performed with this command:

gt ltrharvest -index hopsindex -out hops.out -outinner hops.outinner -gff3 hops.gff > harvScreen.out

Next, LTR_FINDER was run with this command:

ltr_finder -w2 primaryAssembly.fasta > findScreen.out.

Lastly, LTR_retriever was run with this command:

LTR_retriever -genome primaryAssembly.fasta -inharvest harvScreen.out -infinder findScreen.out -threads 16

Repeat masking was performed on the assembly using the library of non-redundant *de novo* hop LTRs with this command:

RepeatMasker -lib hopLTRs.fasta -pa 16 -gff contig.fasta -dir repMask_contigName

Repeat masking was also performed using a database of repeats from MIPS PlantDB using this command: RepeatMasker -lib mipsREdat_9.3p_Eudicot_TEs.fasta -q -xm -small -gff -dir repMask_contigName/ contig.fasta.

The output of running RepeatMasker on the *de novo* LTRs and MIPS PlantDB was combined and used for all downstream analyses.

**Transcriptome assembly**

Transcriptome assemblies from leaf, meristem, and stem were generated by first creating hisat2 indexes with this command:

hisat2-build hopCascade.fasta hopCascade

RNA-seq was aligned to the assembly with hisat2 using this command, which is only shown for leaf tissue:

hisat2 --rna-strandness FR --no-discordant --no-mixed --dta -x HISAT2_INDEXES/hopCascade -1 leaf1_pair1.fastq,leaf2_pair1.fastq -2 leaf1_pair2.fastq,leaf2_pair2.fastq -S hopCascade_vs_leaf.sam.

The sam file was converted to bam using this command:

samtools view hopCascade_vs_leaf.sam -bS -o hopCascade_vs_leaf.bam

Then, the bam file was sorted by coordinates using this command:

samtools sort hopCascade_vs_leaf.bam hopCascade_vs_leaf_coordSorted.bam.

The RNA-seq alignments were assembled into transcripts with stringtie using this command: stringtie hopCascade_vs_leaf_coordSorted.bam -j 2 -o hopCascade_vs_leaf_coordSorted.gtf --fr -A hopCascade_vs_leaf_coordSorted.tab

All three tissue-specific RNA-seq transcriptome assemblies were merged with cuffmerge using this command: cuffmerge -o hopCascade hopCascade.txt.

The ‘hopCascade.txt’ file contains paths to each tissue-specific transcriptome assembly.

**Protein-coding transcript prediction**

Protein coding transcripts were generated from transcriptome assemblies using Transdecoder. The first step was to extract transcripts from the assembly based on coordinates from the merged transcriptome assembly GTF file using this command:

TransDecoder-3.0.1/util/cufflinks_gtf_genome_to_cdna_fasta.pl merged.gtf hopCascade.fasta > hopCascadeTranscripts.fasta.

The next step required getting the longest open reading frame (ORF) with this command:

TransDecoder-3.0.1/TransDecoder.LongOrfs -t hopCascadeTranscripts.fasta

Homology to Embryophyta UniProt genes and Pfam domains was assessed with blastp and hmmscan. Alignment to Embryophyta UniProt genes was performed with this command:

blastp -query longest_orfs.pep –db embryophytaUniprot.fasta -outfmt 6 -evalue 1e-5 -num_threads 10 > hopCascade.outfmt6

Protein domain homology was assessed with this command:

hmmscan --cpu 8 --domtblout hopCascade.domtblout Pfam-A.hmm longest_orfs.pep

For Transdecoder training, the longest ORF CDS from *A. thaliana* TAIR10 was obtained with this command: TransDecoder-3.0.1/TransDecoder.LongOrfs -t Arabidopsis_thaliana.TAIR10.cds.all.fa

Protein-coding transcript prediction was performed using this command:

TransDecoder-3.0.1/TransDecoder.Predict -t hopCascadeTranscripts.fasta --retain_pfam_hits hopCascade.domtblout --retain_blastp_hits hopCascade.outfmt6 --train arabidopsisLongestORFs.cds.fasta.

Only the longest ORF peptides and CDS were retained.

**Hint files for gene prediction**

Gene prediction hint files were created from RNA-seq, predicted protein-coding transcripts from Transdecoder, and alignments to Embryophyta UniProt genes and hop-specific ESTs. All hints were compiled into contig-specific hint files. The first step in the hint preparation pipeline was to align the RNA-seq, genes, and ESTs to the assembly. A first-pass at detecting homology was performed with blast programs, blastx and megablast. First, a blastx search was performed by aligning each contig to Embryophyta UniProt genes. An example of the blastx command is:

blastx -query contig.fasta -db embryophytaUniprot.fasta -out contig_vs_embryophytaUniprot.txt -evalue 1e-5 -outfmt 6 -max_hsps 1

Next, genes with a hit to a given contig were compiled in contig-specific fasta files and aligned to that contig with exonerate. An example exonerate command is:

exonerate --model protein2genome -q uniprotHits.fasta -t contig.fasta --showtargetgff yes --showalignment no -Q protein -T dna --forcegtag true --maxintron 100000 > contig_vs_uniprotHit.gff

Hop-specific ESTs from NCBI and TrichOME database were aligned using the same parameters. First, each contig was aligned to the ESTs with megablast. An example of this command is:

contig.fasta -d ESTs.fasta -o contig_vs_ESTs.txt -e 1e-5 -F F -m 8 -W 28

Next, contig-specific fasta files containing EST sequences with a hit to a given contig were compiled and aligned to that contig using exonerate. An example of an exonerate command is:

exonerate --model est2genome -q estHits.fasta -t contig.fasta --showtargetgff yes --showalignment no -Q dna -T dna --forcegtag true --percent 70 > contig_vs_estHits.gff

RNA-seq-based hint files for gene prediction were generated by first merging all tissue-specific transcriptome assemblies with this command:

samtools merge -nr mergedHopCascade.bam hopCascade_vs_leaf.bam hopCascade_vs_meristem.bam hopCascade_vs_stem.bam

The merged bam file was then sorted with this command:

samtools sort -m 5000000000 mergedHopCascade.bam mergedHopCascade_coordSorted.bam

Intron hints were generated using bam2hints with this command:

bam2hints --intronsonly --source=W --in= mergedHopCascade_coordSorted.bam --out= mergedHopCascade_coordSorted.intron.hint

To obtain exon hints, the bam file was converted to wig format using this command:

perl bam2wig mergedHopCascade_coordSorted.bam.

Bam2wig also required the indexed bam file in the same directory, which was created using this command: samtools index mergedHopCascade_coordSorted.bam mergedHopCascade_coordSorted.bam.bai.

Next, exon hints were obtained using wig2hints with this command:

cat mergedHopCascade_coordSorted.wig | perl wig2hints.pl --width=10 --margin=10 --minthresh=2 --minscore=4 --prune=0.1 --src=W --type=exonpart --radius=4.5 --pri=4 --strand='.' > mergedHopCascade_coordSorted.exonpart.hint

Hints from Transdecoder protein-coding transcripts were obtained by first aligning the longest, complete ORF CDS to the assembly with megablast using this command:

megablast -i contig.fasta -d hopCascadeLongestCompleteORF.cds.fasta -o contig_vs_transdecoderCDS.txt -e 1e-5 -F F -m 8 -W 28

Contig-specific fasta files containing CDS with a hit to a given contig were compiled and aligned to that contig using exonerate with this command:

exonerate --model est2genome -q transdecoderHits.fasta -t contig.fasta --showtargetgff yes --showalignment no -Q dna -T dna --forcegtag true --maxintron 100000 > contig.gff.

Augustus-compatible hint files were then generated.

**Gene prediction**

Gene prediction was performed on each contig individually with Augustus. An example command is:

augustus --strand=both --genemodel=complete --extrinsicCfgFileaugustus.2.7/config/extrinsic/extrinsic.custom.cfg --species=arabidopsis contig.fasta --gff3=on --hintsfile=contig.hint --outfile=contig.gff

**Analysis of coding sequences**

Associate contigs were aligned to all primary contigs using LASTZ. An example command is:

lastz primaryContig.fasta associateContig.fasta --gfextend --hspthresh=20000 --chain --gapped --output=primaryContig_vs_associateContig_lastzPlusStrand.txt --format=maf+ --inner=10000 --identity=80 --strand=plus

Primary contig gene model coordinates were projected to the corresponding positions in the haplotig alignment blocks, and CDS from primary contig and haplotig alignment blocks were then extracted. CDS alignments were processed with MACSE (Ranwez et al. 2011), using the exportAlignment sub-program. An example exportAlignment command is:

java -jar macse_v2.03.jar -prog exportAlignment -align primary_vs_haplotig.cds.fasta -codonForInternalStop NNN -codonForInternalFS --- -charForRemainingFS - -out_NT primary_vs_haplotig.aligned.cds.fasta -out_AA primary_vs_haplotig.aligned.cds.fasta

**Phylogenetic tree and synteny analysis**

A phylogenetic tree of hop and *C. sativa* was constructed by first aligning hop gene models to UniProt Embryophyta genes in both directions with these commands:

blastp –query hopGeneModels.pep.fasta –db embryophytaUniprot.fasta -evalue 1e-5 -outfmt '6 std qcovs' -out hop_vs_uniprot.txt

blastp –query embryophytaUniprot.fasta –db hopGeneModels.pep.fasta -evalue 1e-5 -outfmt '6 std qcovs' -out uniprot_vs_hop.txt

Gene models from *C. sativa* were also aligned to UniProt Embryophyta genes in both directions using these commands:

blastp –query cs10.protein_coding.2905.PEP.fasta –db embryophytaUniprot.fasta -evalue 1e-5 -outfmt '6 std qcovs' -out cannabis_vs_uniprot.txt

blastp –query embryophytaUniprot.fasta –db cs10.protein_coding.2905.PEP.fasta -evalue 1e-5 -outfmt '6 std qcovs' -out uniprot_vs_cannabis.txt

Significant hits to CBDAS and CBDAS-like, BBE-like, and THCAS and THCAS-like genes were extracted for both hop and *C. sativa* with these commands:

python getHopGeneModelSeqs.py hopGeneModels.pep.fasta hop_vs_uniprot.txt uniprot_vs_hop.txt uniprotEmbryophyta.fasta

python getCannabisGeneModelSeqs.py cannabisGeneModels.pep.fasta cannabis_vs_uniprot.txt uniprot_vs_cannabis.txt uniprotEmbryophyta.fasta

The significant hit sequences from hop and *C. sativa* were combined and aligned with clustalw2 using this command: clustalw2 -infile=allHits.fasta -type=PROTEIN -outfile=allHits.aln > allHits.clustalw2.out

Gaps were trimmed from alignments with trimal: trimal -in allHits.aln -gappyout > allHits_noGaps.aln

The file ‘allHits_noGaps.aln’ was converted to phylip-relaxed. Phyml was run with this command:

phyml -i allHits_noGaps.phy -d aa -m Blosum62 -c 4 -a e

The above command produced this file: allHits_noGaps.phy_phyml_tree.txt. The tree file was visualized with ETE3.

MBHs were identified by first aligning significant hits from hop and cannabis to CBDAS, CBDAS-like, THCA, THCA-like, and BBE-like genes using these commands:

blastp -query hopHits.fasta -db cannabisHits.fasta -evalue 1e-5 -outfmt '6 std qcovs' -out hop_vs_cannabis.txt

blastp -query cannabisHits.fasta -db hopHits.fasta -evalue 1e-5 -outfmt '6 std qcovs' -out cannabis_vs_hop.txt.

Regions of synteny were investigated and visualized with Integrative Genomics Viewer (IGV).

Additional custom scripts and details about how they were run can be found at:

<https://github.com/padgittl/CascadeHopAssembly/>

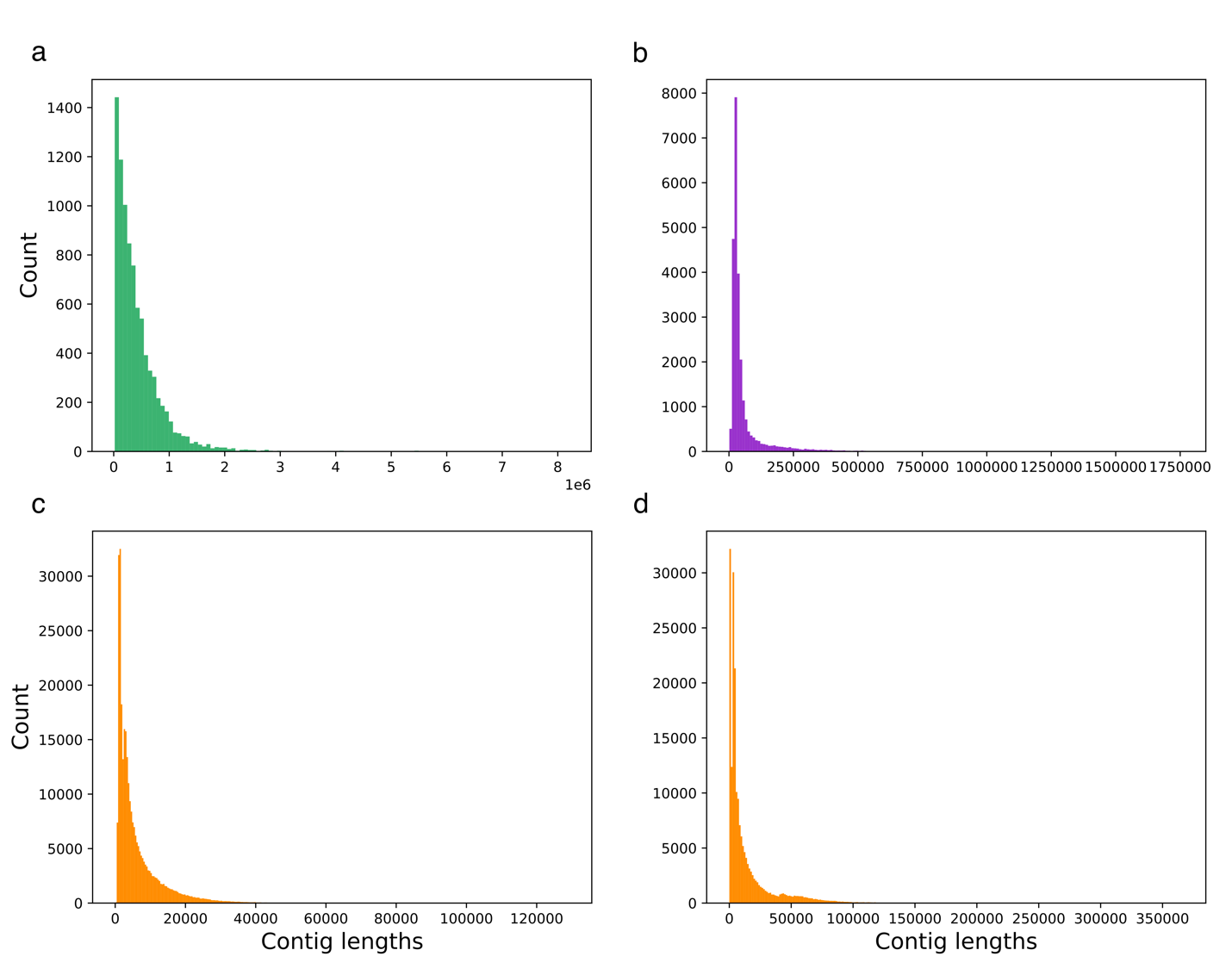

Supplementary Figure 1. (a) Length distribution of primary contigs from the final, deduplicated primary assembly. (b) Length distribution of contigs from the associate assembly. (c) Length distribution of scaffolds from Shinshuwase assembly. (d) Length distribution of scaffolds from Teamaker assembly.

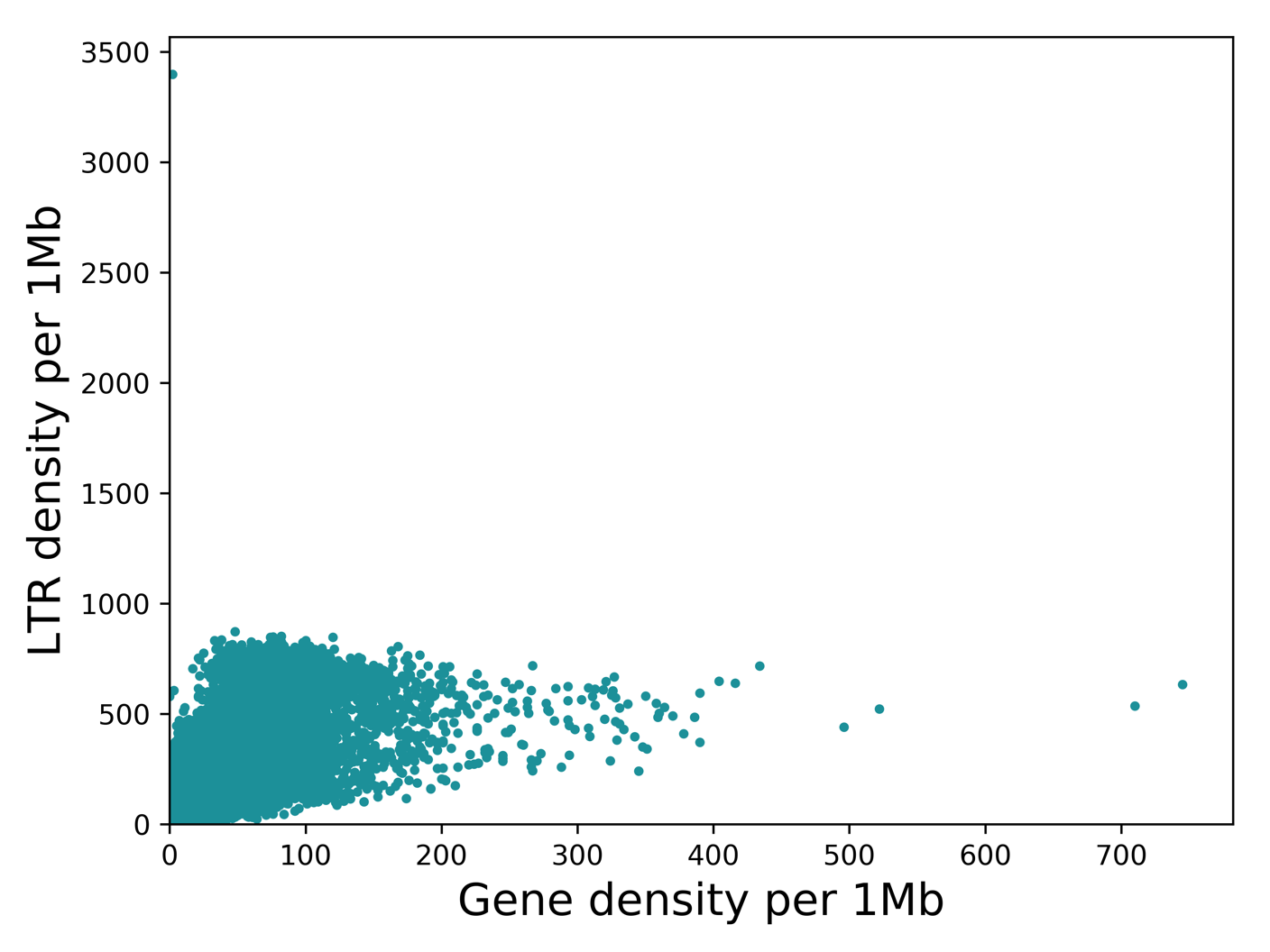

Supplementary Figure 2. Gene and LTR density (Pearson correlation 0.60, p-value 0.0).

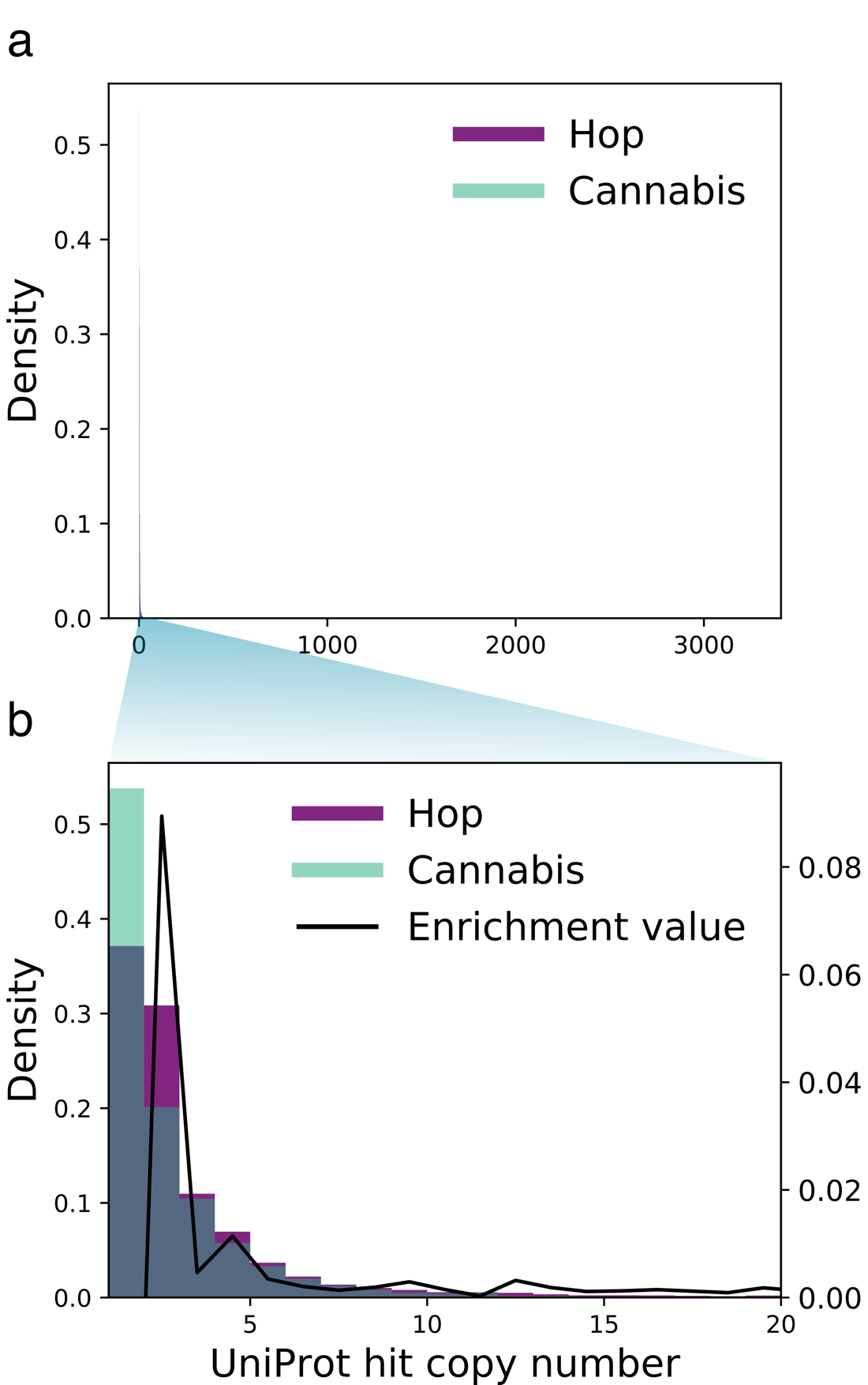

Supplementary Figure 3. Gene copy number density in hop and *C. sativa*. A zoom-in of the region corresponding to 1-20 gene copies features an enrichment spike for hop at two and four gene copies.

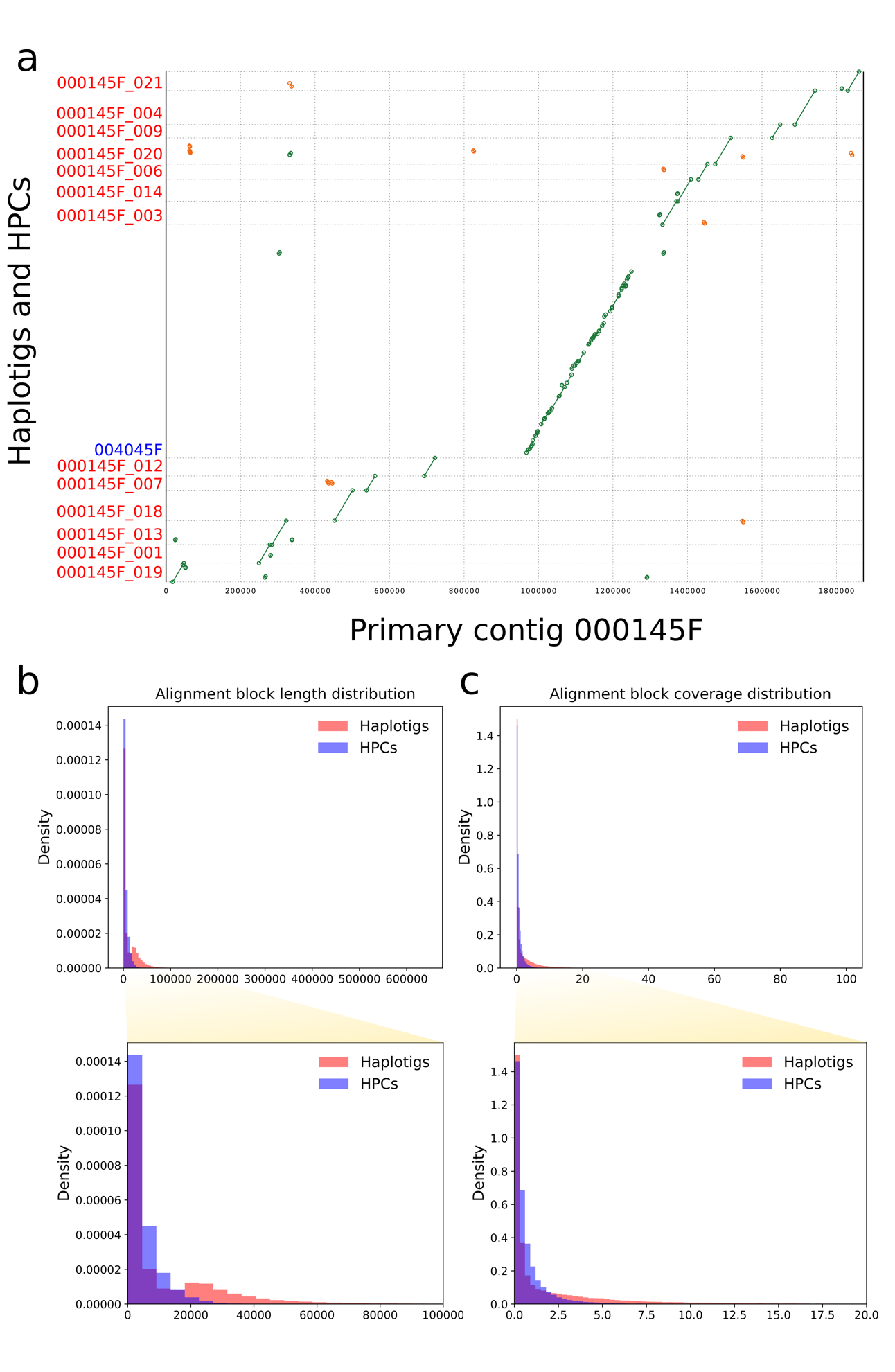

Supplementary Figure 4. Large-scale structural variation assessed by alignment in haplotigs and HPCs. (a) Alignment of a primary contig and its associate contigs demonstrates that haplotig alignments tend to be shorter and have greater homology than alignments to HPCs. (b) A distribution of alignment block lengths shows that haplotig alignments are more abundant at longer lengths than HPC alignments, which are most abundant at shorter alignment block lengths. (c) A distribution of alignment block coverage shows that haplotig alignments are more abundant at longer coverage regions than HPC alignment blocks lengths, and are most abundant at short coverage lengths. HPC coverage is most abundant at intermediate coverage lengths.

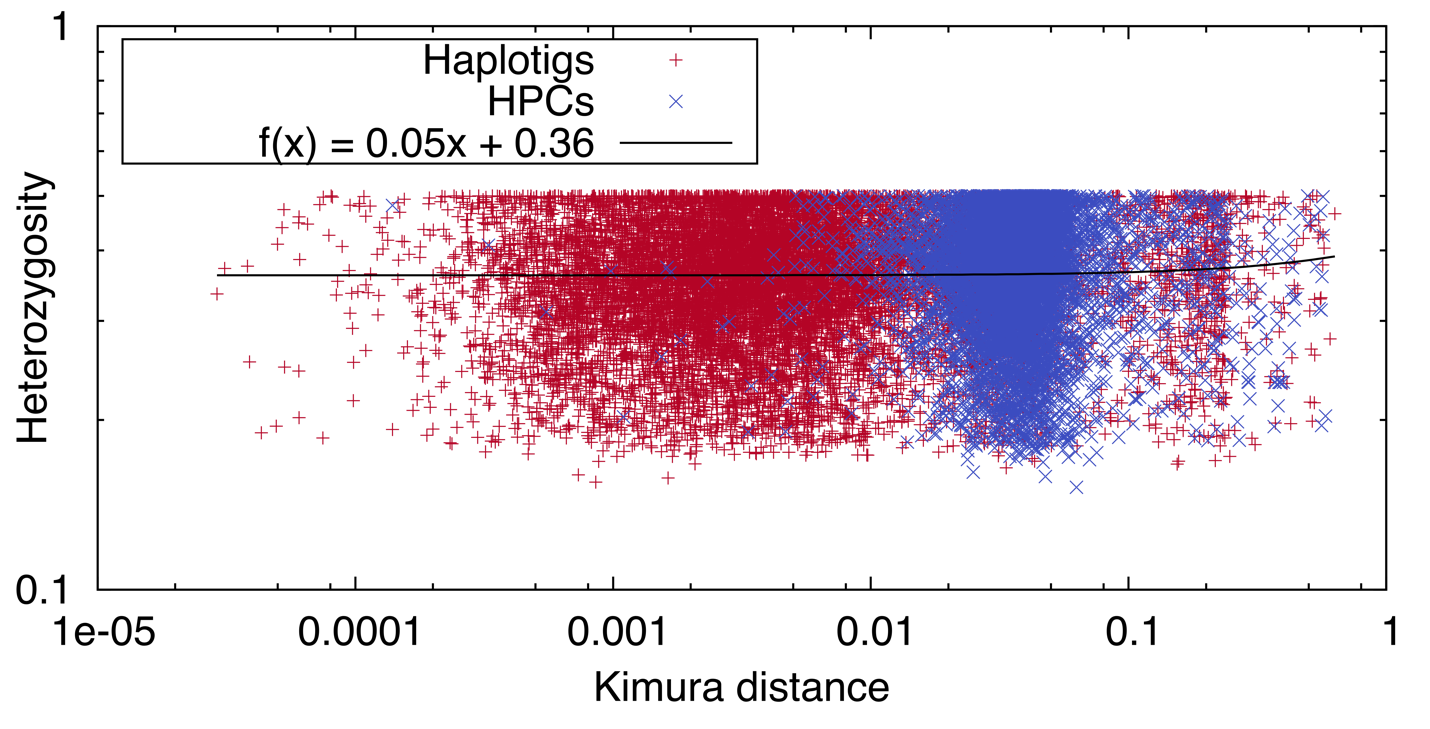
Supplementary Figure 5. Heterozygosity values from primary contigs overlapping haplotigs or HPCs and Kimura distances from the alignment blocks of primary contigs and haplotigs or HPCs.

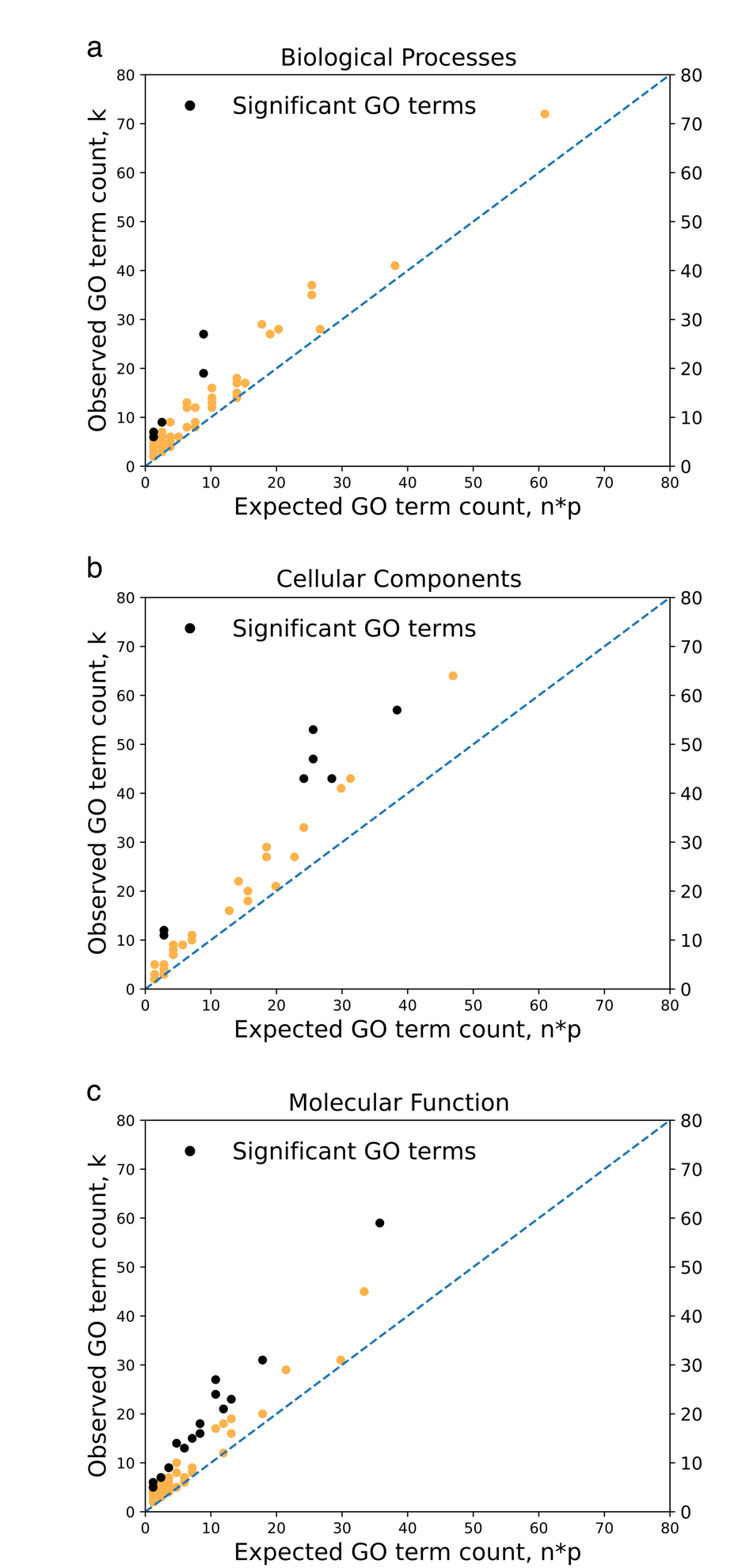

Supplementary Figure 6. The lowest Kimura distance for the top 25% of haplotig genes was used to set the Kimura distance threshold for HPC genes. The value *n* is the number of HPC genes with a Kimura distance above the threshold set by the top 25% of haplotig genes. Probability *p* was obtained from the haplotigs, and is the probability of a GO term occurring in the set of genes in the top 25% of Kimura distances. The observed value *k* is the observed number of occurrences of a GO term in the set of HPCs with the top 25% of Kimura distances. (a) Biological Processes; (b) Cellular components; (c) Molecular function.
